# Suppression of systemic T cell immunity to viral infection during liver injury is prevented by inhibition of interferon and IL-10 signaling

**DOI:** 10.1101/2022.07.21.501031

**Authors:** Carl-Philipp Hackstein, Jasper Spitzer, Konstantinos Symeonidis, Helena Horvatic, Tanja Bedke, Babett Steglich, Lisa M. Assmus, Alexandru Odainic, Nina Kessler, Sabine Klein, Marc Beyer, Ricarda Schmithausen, Eicke Latz, Christian Kurts, Jonel Trebicka, Richard A. Flavell, Natalio Garbi, Axel Roers, Samuel Huber, Susanne V. Schmidt, Percy A. Knolle, Zeinab Abdullah

**Affiliations:** Institute of Molecular Medicine and Experimental Immunology, University Hospital Bonn, Germany; Institute of Innate Immunity, University Hospital Bonn, Germany; Medizinische Klinik und Poliklinik, Hamburg Center for Translational Immunology (HCTI), Universitätsklinikum Hamburg-Eppendorf, 20246 Hamburg, Germany; Medizinische Klinik 1. Universitätsklinikum Frankfurt. Goethe Universität I. Medizinische Klinik und Poliklinik; Molecular Immunology in Neurodegeneration, German Center for Neurodegenerative Diseases (DZNE), Bonn, Germany; Institute for Hygiene and Public Health, University Hospital Bonn, Germany; Department of Immunobiology, Yale University School of Medicine, New Haven, CT, USA; Institute of Immunology, University of Heidelberg, Germany; Institute of Molecular Immunology and Experimental Oncology, Technical University of Munich, Germany; German Center for Infection Research, Munich site

## Abstract

Patients with liver injury such as cirrhosis are at increased risk of intractable viral infections and are hyporesponsive to vaccination. Here, we report that liver injury leads to inhibition of systemic T cell immunity (*LIST*), which abrogated anti-viral immunity and caused persistent infection in preclinical liver injury models. Enhanced gut microbial-translocation but not dysbiosis induced tonic type-I-interferon (IFN) signaling in hepatic myeloid cells, which was responsible for their excessive production of IL-10 after viral infection. Antibiotic treatment reducing intestinal microbial burden or inhibition of IFN- and IL-10-signaling all restored anti-viral immunity without immune pathology. Importantly, inhibition of IL-10 restored virus-specific immune responses to vaccination in cirrhotic patients. Thus, *LIST* results from sequential events involving intestinal microbial translocation, hepatic myeloid cell-derived IFN-/IL-10 expression, and finally inhibitory IL-10 receptor-signaling in T cells, of which IL-10Rα-signaling may serve as target to reconstitute anti-viral T cell immunity in cirrhotic patients.

## Main

Patients with severe liver damage and liver cirrhosis respond poorly to vaccination^1,2^, and often suffer from severe complications during influenza virus infection including organ failure, secondary infections, and death^3-5^. Vaccine hypo-responsiveness and susceptibility to infection in patients with severe liver injury were suggested to be due to profound abnormalities in B cell phenotype and function triggered by TLR-mediated sensing of translocated microbiota^6^. However, the mechanisms underlying impaired adaptive immune responses remain poorly understood, and were mostly limited to studies of LPS challenge as surrogate model for bacterial infection^7^. In the last years, the prevalence of non-alcoholic fatty liver disease and alcohol-related liver disease leading to liver fibrosis, cirrhosis, and liver cancer is dramatically increasing^8^. In addition to sincere consequences of the loss of functional liver tissue, patients with severe liver injury are more susceptible to bacterial^2^ and viral infections^3,5,9-11^, which are major factors driving patient morbidity and mortality^12^. Further, the efficiency of vaccinations against relevant human pathogens like IAV, HAV and HBV has been reported to be considerably lower in cirrhosis patients^1,4^, leading to an increased risk of potentially life-threatening infection. Taken together, these observations suggest that the ability of the immune system to mount a proper adaptive response is impaired during liver injury.

Dysfunction of T cells resulting from co-inhibitory signaling and/or continuous (hyper-) activation has been described in the context of malignant diseases and chronic viral infection^13^, and is characterized by a progressive loss of T cell functionality as well as overall quantity, ultimately rendering the immune system unable to contain pathogens. Excessive and chronic signaling by type I interferons (IFN I) has been identified as a key factor driving T cell dysfunction in preclinical models of viral infections^14-17^. Interestingly, upregulation of IFN I is also a main feature of cirrhosis^18,19^, and we have previously shown that IFN I, especially in the context of infection, is responsible for impairment of innate immunity in murine and human myeloid immune cells in the liver^18^. Here, we demonstrate impairment of the adaptive arm of the immune system during liver injury through IFN I signaling inducing dysfunction in both CD8 and CD4 T cells leading to loss of antiviral T cell surveillance. Using complementary experimental approaches, we show that tonic IFN I signaling during liver injury resulted in production of high levels of the immune regulatory cytokine IL-10. We identified IL-10 as the key mediator of T cell dysfunction during liver injury and provide evidence that blocking of IL-10 rescues T cell function and allows the control of viral infection without organ immune pathology. Given the importance of CD8 and CD4 T cell responses for immune surveillance our findings contribute to the understanding of failing immune responses in cirrhotic patients and identify the IL-10Rα-signaling pathway as a potential molecular target to improve immune control and vaccination efficacy in these patients.

### Defective immune responses to vaccination in patients with liver cirrhosis

To characterize the reported low vaccine responsiveness in patients with severe liver injury ^1,2^, we compared the responses of cirrhotic patients to those of healthy individuals after vaccination against hepatitis B in a proof-of-concept clinical study (**Extended Data Fig. 1a, Supplementary Table I**). Healthy individuals had high antibody titers against the surface antigen of HBV (anti-HBs) at six months after vaccination, whereas anti-HBs antibody titers in cirrhotic patients were much lower and often below the cutoff level of 100 U/l (**Fig. 1a**) that is considered necessary for protection against infection^20^. Furthermore, peripheral blood-derived HBs-specific CD8 and CD4 T cells from cirrhotic patients failed to proliferate and to produce the effector cytokines IFNγ and IL-21 after *ex vivo* stimulation with HBs-specific peptides^21^ (**Fig. 1b-e, Supplemental Table II**). Additionally, we assessed the antibody response to COVID-19 vaccination in infection naïve cirrhotic patients or age- and sex-matched healthy individuals (**Supplementary Table III**). All individuals received two doses of the COVID-19 mRNA (BNT162b2) vaccine, with the second vaccine dose being applied on average 32 days after the first dose, and blood samples for analysis were obtained before and 7-10 days after the second vaccination (**Extended Data Fig. 1b**). Clearly, COVID-19 vaccine-induced antibody titers against the SARS-CoV-2 spike protein were significantly lower in cirrhotic patients compared to healthy individuals (**Fig. 1f**). Of note, vaccination-induced anti-RBD IgG binding to SARS-CoV-2 variants of concern such as B1.351 (beta), B.1.1.7(alpha) and P.1(gamma) was significantly lower in cirrhotic patients (**Fig. 1g**). Together these results indicate poor development of virus-specific B and T cell immunity after vaccination in cirrhotic patients.

**Figure 1.**
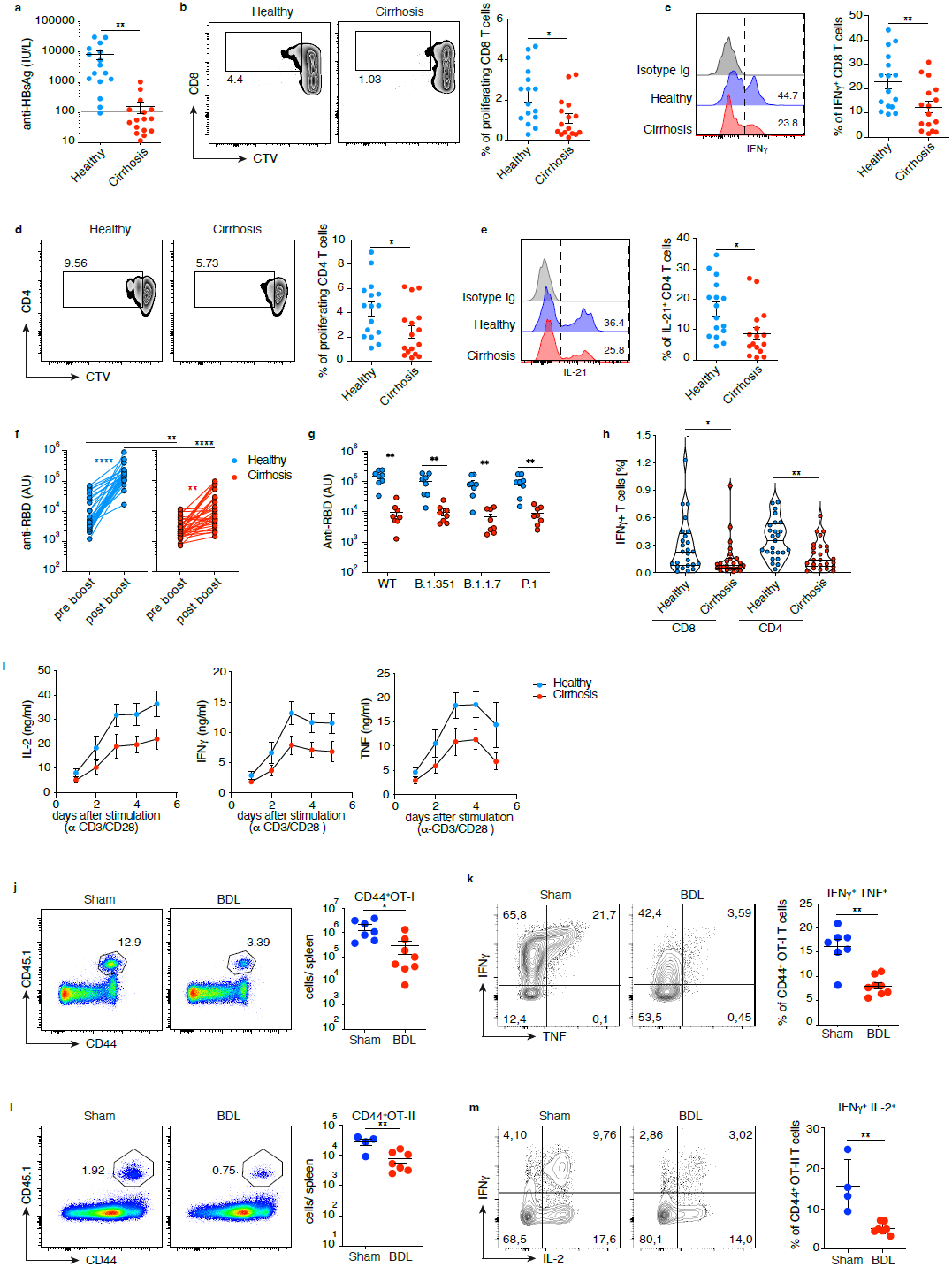
Poor immune responses to vaccination in cirrhotic patients and mice with liver injury. **a**, serum anti-HBs IgG titres in sera of healthy donors (n=16) and cirrhotic patients (n=16) after completion of a full set of vaccinations against HBsAg. **b-e**, frequencies of proliferating and IFNγ-producing CD8 T cells or IL-21-producing CD4 T cells from healthy donors or cirrhotic patients 6 months after third vaccination and after *ex vivo* stimulation with HBsAg peptides. **f**,**g**, anti-SARS-CoV-2 spike serum IgG (arbitrary units) in healthy donors or cirrhotic patients at d7-10 after the second COVID-19 mRNA vaccination (BNT162b2), and scatter plot depicting serum IgG binding (AU) to the RBD of SARS-CoV-2 VoCs. **h**, frequencies of IFNγ-producing CD8 and CD4 T cells after *in vitro* stimulation with peptides covering the SARS-CoV-2 spike protein from healthy subjects or cirrhosis patients d7-10 after booster vaccination. **i**, IL-2, IFNγ- and TNF-release from anti-CD3/CD28 stimulated T cells from healthy individuals or cirrhotic patients. **j-m**, frequencies and *ex vivo* cytokine production of CD45.1^+^ OT-I (**j**,**k**) and OT-II T cells (**l**,**m**) previously transferred into BDL- or sham-operated mice that were vaccinated with ovalbumin and pI:C (data shown for d7 post vaccination). **i-m** data representative of three independent experiments. **a-e**,**g**,**h**,**i-m** statistical analysis by unpaired t-test, *P≤0.05, **P≤0.01, **f**, paired Wilcoxon **P≤0.01, ****P≤0.0001.

Importantly, the failure of cirrhotic patients to mount strong T cell responses was not restricted to vaccination. Polyclonal stimulation of circulating T cells from cirrhotic patients with anti-CD3/CD28 antibodies was associated with lower production of cytokines like IL-2, IFNγ and TNF (**Fig. 1h**). Compared to healthy controls, CD8 T cells from cirrhotic patients were more likely to undergo apoptosis indicated by higher numbers of AnnexinV^positive^ cells (**Extended Data Fig. 1d**), had lower mitochondrial membrane potential and higher numbers of depolarized mitochondria (**Extended Data Fig. 1e,f**), and higher baseline level of reactive-oxygen species that failed to increase upon stimulation (**Extended Data Fig. 1g**). Collectively, these findings indicated an inhibitory mechanism broadly hampering adaptive immunity in cirrhotic patients. To study this inhibition of T cell immunity after vaccination *in vivo*, we employed a preclinical model of severe liver injury, i.e., bile duct ligation (BDL) that induces liver fibrosis and cirrhosis^18^. To monitor T cell immunity after vaccination, we adoptively transferred naïve ovalbumin-specific CD45.1^+^CD8 (OT-I) or CD4 (OT-II) transgenic T cells one day prior to s.c. vaccination with ovalbumin and the TLR3-ligand polyI:C as adjuvant. On day 8 post vaccination, significantly lower numbers of splenic antigen-specific CD44^hi^ CD45.1^+^CD8 and CD4 T cells were found in BDL-compared to sham-operated mice (**Fig. 1i,k**) suggesting reduced T cell activation and expansion. Furthermore, CD45.1^+^CD8 and CD4 T cells from BDL-mice produced lower levels of the effector cytokines IFNγ, TNF and IL-2 upon *ex-vivo* restimulation (**Fig. 1j,l**). Taken together, these findings in BDL-mice confirmed dysfunctional T cell response to vaccination in cirrhotic patients and led us to characterize the molecular mechanisms underlying liver injury-associated loss of T cell immunity.

### Failure of anti-viral T cell immunity to control viral infection in mice with liver injury

To explore the impact of severe liver injury on antiviral T cell immune surveillance, we used mice with liver injury from two different etiologies, i.e., BDL or repeated application of carbon tetrachloride (CCl_4_), for a challenge through infection with the lymphocytic choriomeningitis virus (LCMV-WE and LCMV-Armstrong strain) that is rapidly cleared by the immune system of healthy mice^22,23^. In BDL-mice, LCMV replication peaked at slightly higher viral titers but with similar time kinetics in blood, liver and spleen compared to sham-treated mice (**Fig. 2a-c**). Strikingly, BDL-mice failed to clear LCMV-WE from blood, liver and spleen and LCMV replication persisted for at least 30 days post infection, whereas sham-treated mice successfully controlled LCVM replication latest within 12 days (**Fig. 2a-c, data not shown**).

**Figure 2.**
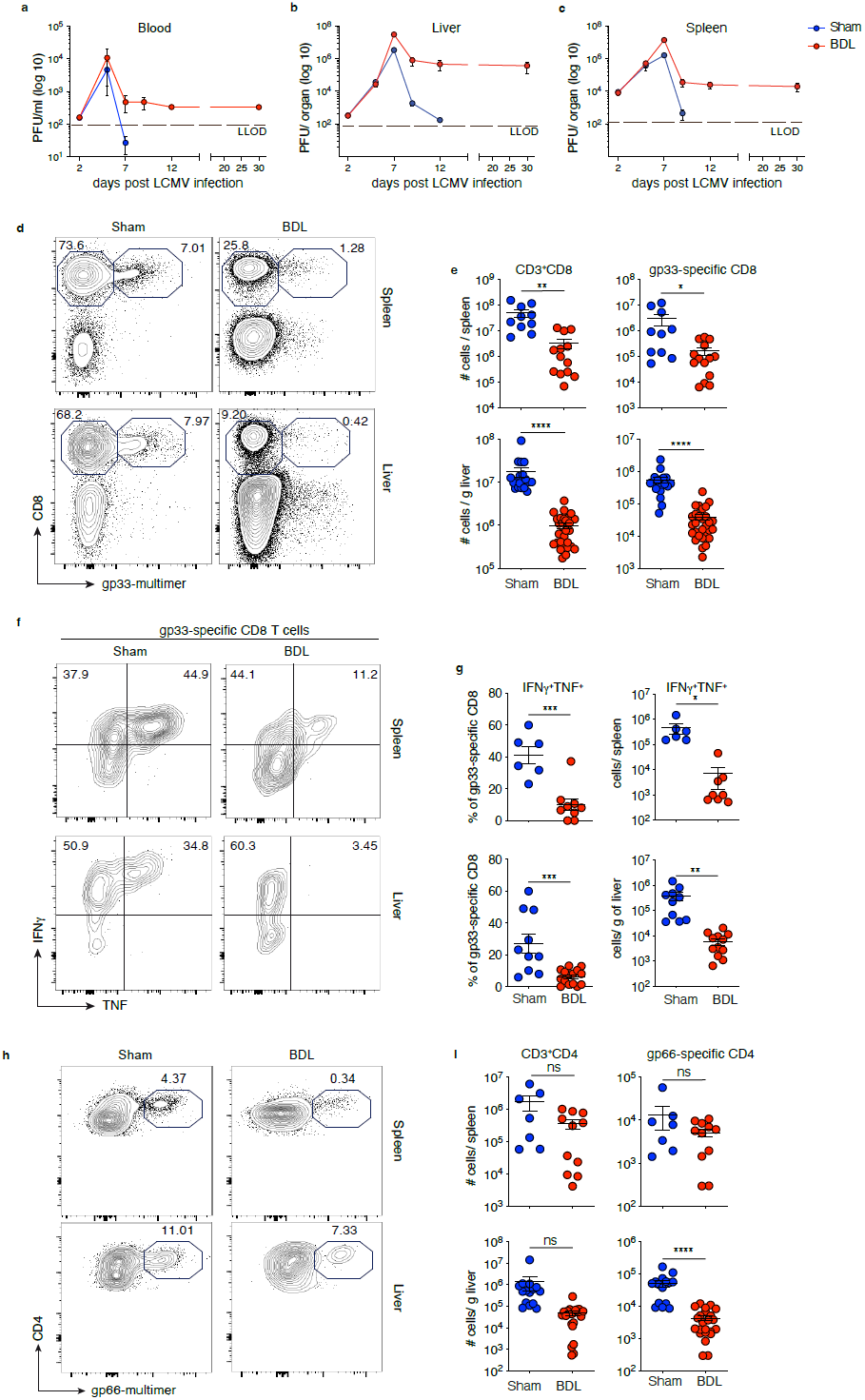
Liver injury in mice is associated with failure to clear LCMV infection and impaired antiviral T cell responses. **a-c**, dynamics of LCMV titers determined as plaque-forming units (PFU) in blood, liver and spleen of BDL- or sham-operated mice after LCMV infection (2×10^4^ PFU) (n=5). LLOD – lower limit of detection. **d**,**e**, quantification of total CD8 T cells and gp33-specific CD8 T cells from spleen and liver of BDL- or sham-operated mice d8 p.i. **f**,**g**, IFNγ and TNF production by spleen- or liver-derived LCMV gp33-specific CD8 T cells after *ex vivo* stimulation with PMA/Ionomycin and quantification. **h**,**i**, frequencies of CD4 T cells and LCMV gp66-specific CD4 T cells isolated from spleen and liver of BDL-or sham-operated mice at d8 p.i. **a**-**c**, data representative of ≥3independent experiments; **e**,**f**,**i**, pooled data from ≥ 3 independent experiments; statistical analysis by unpaired t-test; ns = not significant, *P≤0.05, **P≤0.01, ***P≤0.001, ****P≤0.0001.

LCMV persistence also in the spleen indicated a systemic loss of antiviral immune surveillance, rather than a local attenuation of anti-viral immunity selectively in injured liver tissue. This fundamentally different course of viral infection prompted us to determine virus-specific T cell immunity at the peak of effector response, i.e. day 8 post infection^23-25^, when failure to control viral replication became apparent in BDL-mice. Consistent with efficient control of LCMV infection, sham-treated mice displayed strong LCMV-specific CD8 T cell immunity, quantified by dextramer-positive CD8 T cells recognizing the gp33 LCMV epitope. BDL-mice, in contrast, had 50 to 100-fold lower numbers of LCMV-specific CD8 T cells (**Fig. 2d,e**, Extended Data Fig. 2d,e). Moreover, most LCMV gp33-specific CD8 T cells detected in BLD-mice were dysfunctional with reduced IFNγ and TNF expression after peptide-specific restimulation, which together with reduced numbers led to a >100 fold reduction in TNF-producing LCMV-specific CD8 T cells in BDL-mice (**Fig. 2f,g**). Similar persistence of LCMV infection and loss of virus-specific CD8 T cell immunity was detected in mice with liver injury after repeated CCl_4_-application (**Extended Data Fig. 2a-g**), indicating that liver injury independent of its cause led to loss of anti-viral immune surveillance.

To study defects in virus-specific CD8 T cells in more detail, we transferred TCR-transgenic P14 CD8 T cells, that express a T cell receptor specific for the LCMV-gp33 epitope, into BDL-mice the day before LCMV infection. LCMV infection in BLD-mice showed 100 to 1000-fold reduced numbers of LCMV-specific P14 CD8 T cells compared to healthy mice and the few P14 T cells present were dysfunctional with reduced expression of IFNγ/TNF (**Extended Data Fig. 2h-k**), confirming the results obtained from studying virus-specific T cells generated from the endogenous TCR repertoire. Of note, we also detected reduced numbers of LCMV-specific CD4 T cells in BDL-mice (**Fig. 2h,i**), that had diminished effector cytokine production with only few cells co-expressing IFNγ, TNF and IL-2 after peptide-specific stimulation (**Extended Data Fig. l**,**m**). Together, these experiments demonstrated a broad and severe dysfunction of virus-specific CD8 and CD4 T cell immunity in mice with liver injury that was associated with failure to control LCMV infection.

### Virus specific T cells during liver injury rapidly become dysfunctional after infection

To map the landscape of transcriptional changes that distinguishes antigen specific CD8 T cells during liver injury, we performed RNA-seq analysis of sorted P14 cells from BDL- or sham-operated mice at day 8 post LCMV infection. This identified 2153 differentially regulated genes in dysfunctional P14 T cells (**Extended Data Fig. 3a, Supplementary Table III**). Gene Ontology enrichment analysis (GOEA) and network visualization revealed differences in biological processes associated with lymphocyte activation, immune effector functions, metabolic and signal transduction processes (**Fig. 3a**). Genes encoding inhibitory receptors (*Cd160, Havrc2, Pdcd1, Ctla4, 2b4*, and *Lag3*) and inflammation-associated cytokines like *Tnf* and *Il10* were up-regulated in T cells from BDL-mice, whereas transcription factors associated with effector T cell functions^26^ namely, *Tbet, Tcf7, Eomes* and *Bcl6* were down-regulated (**Fig. 3b,c**). Dysfunctional T cells were further characterized by increased expression of transcription factors associated with previously T cell exhaustion such as *Tox, Batf, Irf4* and *Id3*^*27-29*^ (**Fig. 3c**). To test whether the gene signature of dysfunctional T cells from mice with liver injury bears similarities with the T cell exhaustion signature observed in cancer and chronic viral infections, we performed gene set enrichment analysis (GSEA) using an established gene signature of T cell exhaustion^27^. P14 cells from BDL-mice showed a significant enrichment for genes upregulated in exhausted T cells (**Fig. 3d**), and featured hallmarks of the transcriptomic signature of exhausted CD8 T cells (**Fig. 3e**). Flow cytometric analysis confirmed expression of these genes at the protein level in both transferred P14 (**Fig. 3f**) as well as splenic endogenous LCMV-specific CD8 T cells of BDL-mice (**Extended Data Fig. 3b,c**). We observed increased numbers of virus-specific CD8 and CD4 T cells co-expressing PD1, TIM3 and LAG3 in liver and spleen from both BDL-as well as CCl_4_-treated mice (**Extended Data Fig. 3d-f**). Similarly, adoptively transferred P14 T cells in BDL-mice demonstrated high level expression of BATF, TOX and IRF4 expression on day 8 p.i. (**Fig. 3f and Extended Data Fig. 3c**). These data demonstrate that loss of antiviral T cell surveillance develops during liver injury, irrespective of the underlying injury etiology.

**Figure 3.**
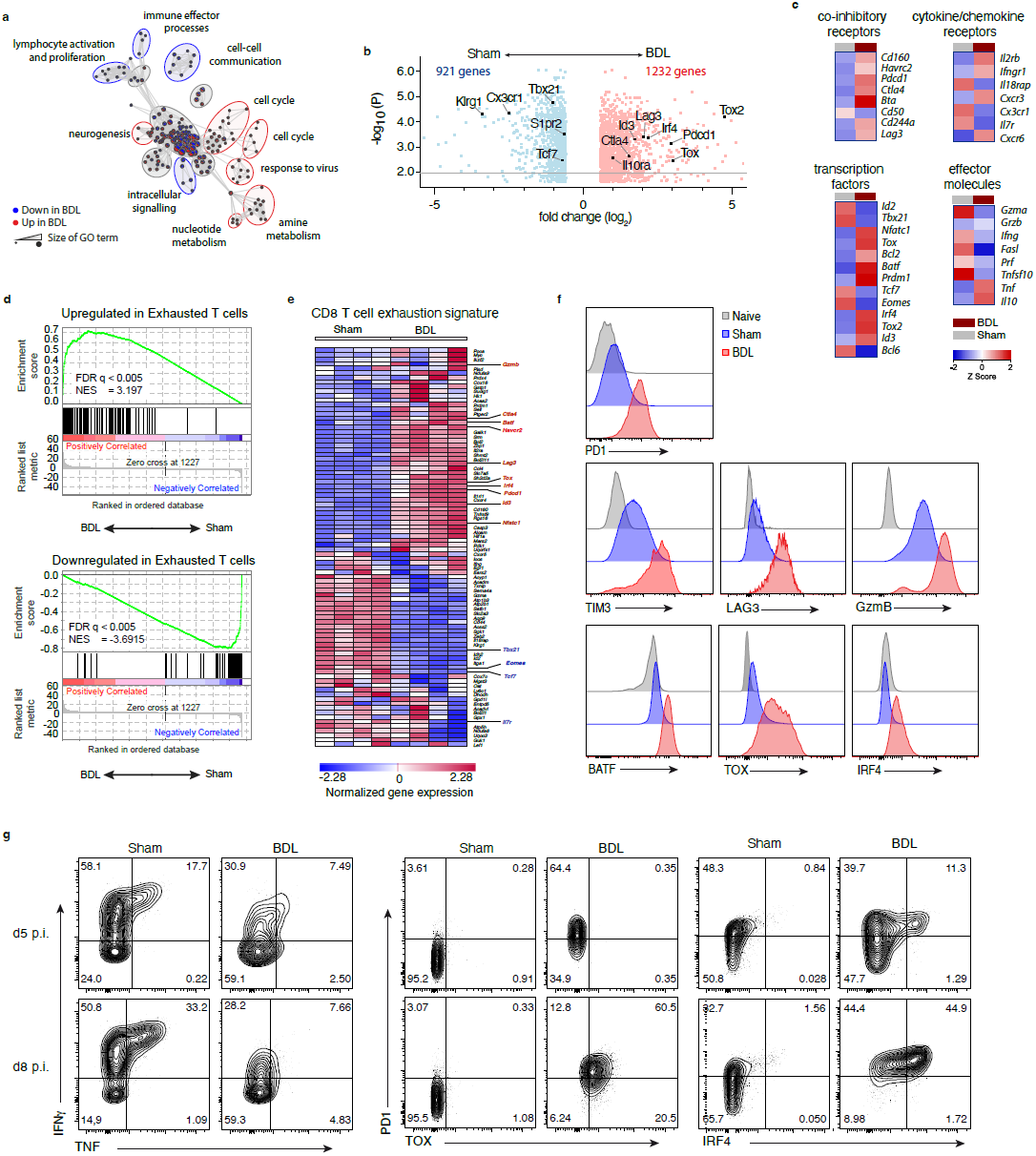
Upregulation of dysfunction-associated molecules in T cells during liver injury. **a**, Bingo network of enriched gene ontology terms for 921 down-regulated (blue) and 1232 up-regulated (red) genes in in LCMV-specific (P14) CD8 T cells from BDL-compared to sham-operated mice at d8 p.i. (LCMV-WE) (n=4). **b**, volcano plot of 2153 differentially expressed genes in P14 T cells from BDL-or sham-operated mice at d8 p.i. **c**, heatmap of centered mean expression of selected genes in P14 CD8 T cells as in (b). **d**, Gene Set Enrichment Analyses (GSEA) using a gene set characterising exhausted T cells during chronic viral infection^27^ for the DEGs from (b). **e**, heatmap visualizing mRNA expression levels of classical signature genes for CD8 T cell exhaustion^27^ in DEGs from (b). **f**,**g**, co-expression levels of co-inhibitory surface receptors and transcription factors in P14 CD8 T cells from BDL- or sham-operated mice at d8 (**f**) or d5/d8 p.i. (**g**). **f**,**g**, data representative of two independent experiments.

We next assessed if T cell dysfunction developing during liver injury coincided with expression of TOX, BATF and IRF4. For this, we evaluated effector function and transcription factor expression in T cells on day 5 p.i, when precursors of exhausted T cells are known to be present^30^. Although P14 T cell dysfunction in BDL-mice was already present on day 5 p.i., no increased expression levels of TOX or BATF were detected and only some IRF4 expressing P14 T cells were found (**Fig. 3g**, data not shown). These results point towards a distinct cause of T cell dysfunction during liver injury compared to TOX-dependent exhausted T cells, and prompted us to identify the underlying mechanisms.

### CD4 and CD8 T cell dysfunction during liver injury associated with IFN I-signaling

During chronic liver injury elevated expression of immune regulatory molecules is found^18,31,32^, which may curtail antiviral T cell immunity. GSEA revealed enrichment of 148 genes associated with TGFβ-signaling in P14 T cells from BDL-mice (**Fig. 4a, Extended Data Fig. 4a**). Consistently, we detected increased levels of TGFβ expression (**Fig. 4b**) in mice with liver injury. Further, LCMV-specific CD8 T cells from BDL-mice showed enhanced levels of phospho-SMAD2 (**Fig. 4c**), a key downstream molecular target of TGFβ receptor signaling^33^. To determine the contribution of TGFβ-receptor signalling for T cell dysfunction we treated BDL-mice with TGFβRII-blocking antibodies the day prior to LCMV infection. Strikingly, almost all BDL-mice succumbed within two days after LCMV infection when TGFβ-receptor signalling was blocked (**Fig. 4d**) without evidence for increased antiviral immune surveillance (**Fig. 4e**), suggesting a critical nonredundant and specific function of TGFβ in tissue-protection but not immune surveillance during liver injury.

**Figure 4.**
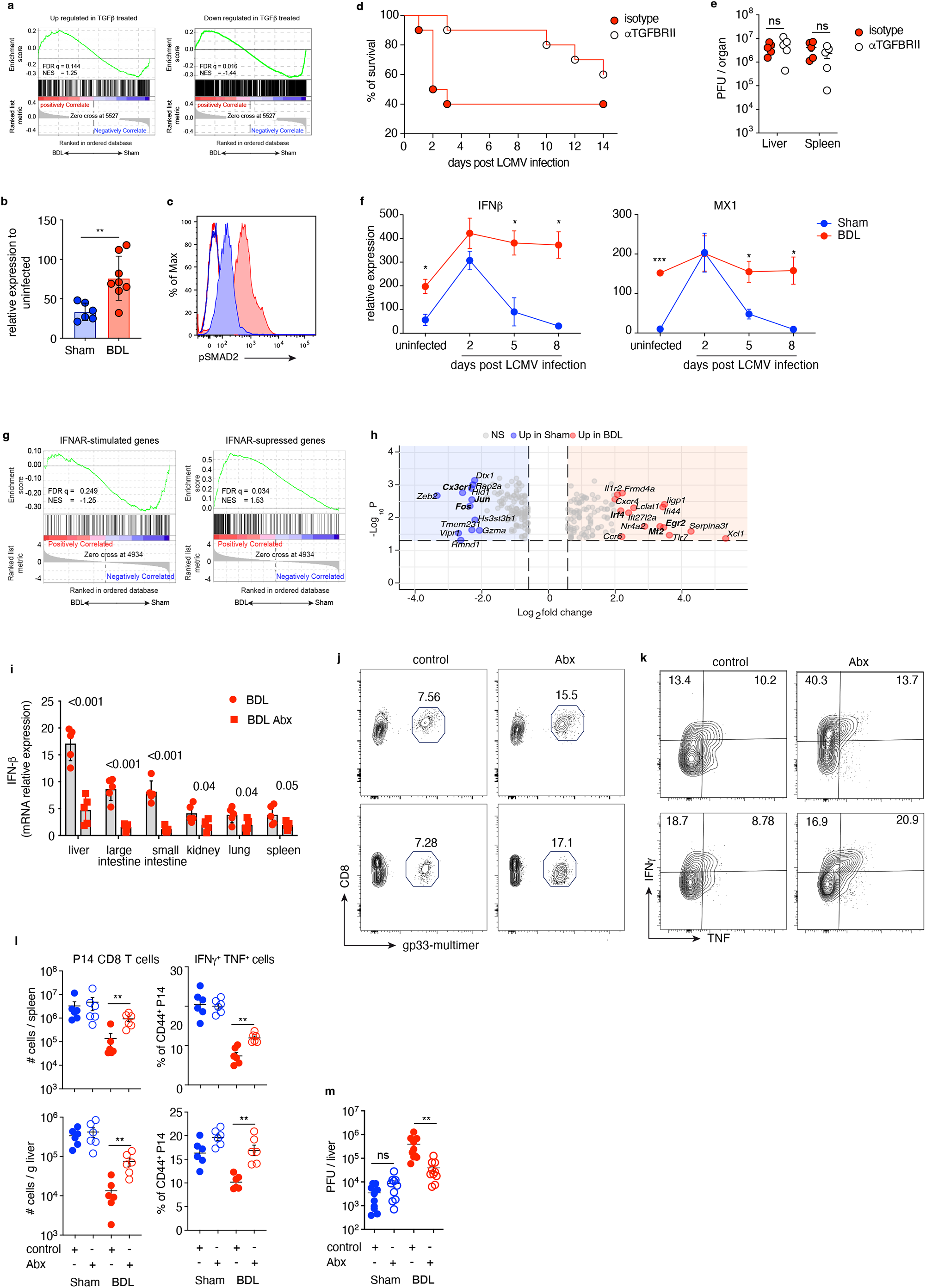
Virus-specific CD8 T cells in BDL-mice are exposed to TGFβ- and IFN I-signaling. **a**, GSEA depicting enrichment of gene sets significantly up-(left) or down-regulated (right) in response to TGFβ-treatment^77^ in the transcriptome of P14 CD8 T cells at d8 p.i. from BDL-mice. **b**, relative expression of TGFβ-expression in the liver of LCMV-infected BDL- or sham-operated mice d8 p.i. **c**, phosphorylated Smad2 in gp33-specific CD8 T cells from spleen of BDL- or sham-operated mice at d8 p.i. **d**, Kaplan-Meier survival curves for LCMV-infected BDL-mice receiving anti-TGFβR-II or isotype control Ig (n=10). **e**, LCMV titers in liver and spleen of mice from (d) on day 8 p.i. **f**, *IFNb* or *Mx1* mRNA expression levels relative to *Hrtp* in liver tissue of BDL- or sham-operated mice after LCMV infection (n=5). **g**, GSEA of gene sets up-(left) or downregulated (right) in response to IFN I-signalling^78^ in cells from (a). **h**, volcano plot analysis of 212 IFN I-related genes at the intersection of differentially expressed genes, top regulated genes with a logFC>|2| in red (up-) and blue (down-regulated). One-way ANOVA according to Benjamini-Hochberg FDR-corrected p-value < 0.05. **i-m**, BDL-mice receiving antibiotics (Abx) in drinking water received P14 cells one day prior to LCVM infection. **i**, IFN I mRNA expression of IFNβ in tissues from BDL-mice (d9 post operation). **j-l**, abundance of P14 T cells and frequency of IFNγ− and TNF-producing P14 T cells in liver and spleen on d8 p.i. **m**, quantification of LCMV titers in liver of mice treated as in (l). **c**,**e**,**f**,**i**,**j**, data from ≥ 2 independent experiment. **c**,**e**,**f**,**i**,**j**,**l** statistics were assessed by unpaired t-test, ns = not significant, *P≤0.05, **P≤0.01, ***P≤0.001.

We also detected high expression of IFN I and the interferon-stimulated gene *Mx1* in mice with liver injury, the expression of which further increased after LCMV infection (**Fig 4f**). While in sham-operated infected mice *Ifnb* and *Mx1* expression declined to base-line on d4 p.i., in mice with liver injury *Ifnb* and *Mx1* continued to increase after infection (**Fig 4f**). GSEA indicated a significant enrichment of 230 genes associated with IFNAR-signaling (from the Interferome data base^34^) in P14 T cells from BDL-mice (**Fig. 4g, Extended Data Fig. 4b**). Among the top 20 upregulated genes from this group, we found *Irf4, Nr4a2, Mt2, Egr2, Lclat1* and *Frmd4a* (**Fig. 4h**), genes which are associated with impaired T cell effector functions in cancer and chronic viral infections^27,35-38^. In contrast, transcription factors involved in signaling like *Jun* and *Fos* were amongst the most down-regulated genes (**Fig. 4h**), altogether indicating a potential key role of IFN I in induction of T cell dysfunction.

Enhanced translocation of gut microbiota (**Extended Data Fig. 4c,d**) and consequently increased innate immune sensing result in tonic type I interferon alpha receptor (IFNAR) signaling in myeloid cells^18^ during liver injury in mouse and man (**Extended Data Fig. 4e,f**). Of note, reduction of intestinal microbial burden by antibiotics treatment led to significant reduction of IFN I expression in the liver and intestine of BDL-operated mice (**Fig. 4i**), improved expansion of effector cytokine production by LCMV-specific CD8 T cells (**Fig. 4j-l**) and enhanced viral clearance (**Fig. 4m)**. Interestingly, colonization of germ-free mice with the microbiome of sham- or BDL-operated mice led to similar levels of IFN I during liver injury indicating that IFN I expression was independent of the gut microbiota composition (**Extended Data Fig. 4g**). These results demonstrated a critical role of microbiota-induced IFN I in T cell dysfunction during liver injury.

### High IFN I expression during liver injury drives loss of T cell immunity

To investigate the role of IFN I-signaling in the loss of T cell immunity during liver injury, we applied an IFNAR-blocking antibody. Inhibition of IFNAR-signaling (**Extended Data Fig. 5a**) shortly after LCMV-infection prevented generation of virus-specific CD8 T cells (data not shown), consistent with the reported key role of IFN I for T cell priming and expansion^39,40^. Strikingly, inhibition of IFNAR-signaling from d5 p.i. onwards increased the numbers of LCMV-specific CD8 and CD4 T cells in mice with BDL-as well as CCl_4_-induced liver injury (**Fig. 5a,b Extended Data Fig. 5b,c,e,g,h**). Blockade of IFN I further rescued from loss of T cell effector function in mice with liver injury, in both BDL and CCl_4_ models of liver injury, shown by increased expression levels of IFNγ and TNF after stimulation (**Fig. 5a,b**, **Extended Data Fig. 5d,f,g,h**). Consequently, we observed a pronounced reduction in the viral load in anti-IFNAR-treated mice with liver injury (**Fig. 5c,d**), almost to the same level as in mice without liver injury. This indicated that elevated levels of IFN I in mice with liver injury were involved in loss of T cell immunity and impaired viral clearance, consistent with the role of IFN I in persistence of LCMV clone 13 and HIV infection^15,16,41^. LCMV-specific CD8 T cells after blockade of IFN I expressed lover levels of co-inhibitory receptors PD1, TIM3 and LAG3 (**Fig. 5e, Extended Data Fig. 5i**) and the transcription factors TOX, IRF4 and BATF (**Fig. 5e,j**). Of note, blocking IFNAR in mice with liver injury induced increased numbers of TCF1^+^P14 T cells and TCF1^+^TIM3^-^ progenitor-exhausted cells (**Fig. 5e, Extended Data Fig. 5j**). To explore whether IFNAR-signaling specifically in T cells might be responsible for their dysfunction during liver injury, we generated CD4-Cre^*ERT2*^ *x* IFNAR^fl/fl^ mice, in which tamoxifen injection deleted IFNAR selectively in ∼60% of T cells (**Extended Data Fig. 6a**). Tamoxifen application from day 3 after LCMV infection did not restore T cell function (**Extended Data Fig. 6b,c**) and did not improve control of LCMV infection (**Extended Data Fig. 6d**). Although this does not rule out a direct effect of IFN I signaling on LCMV-specific CD8 T cells, it rather suggests a role of IFNAR-signaling on other immune cell populations that then causes T cell dysfunction.

**Figure 5.**
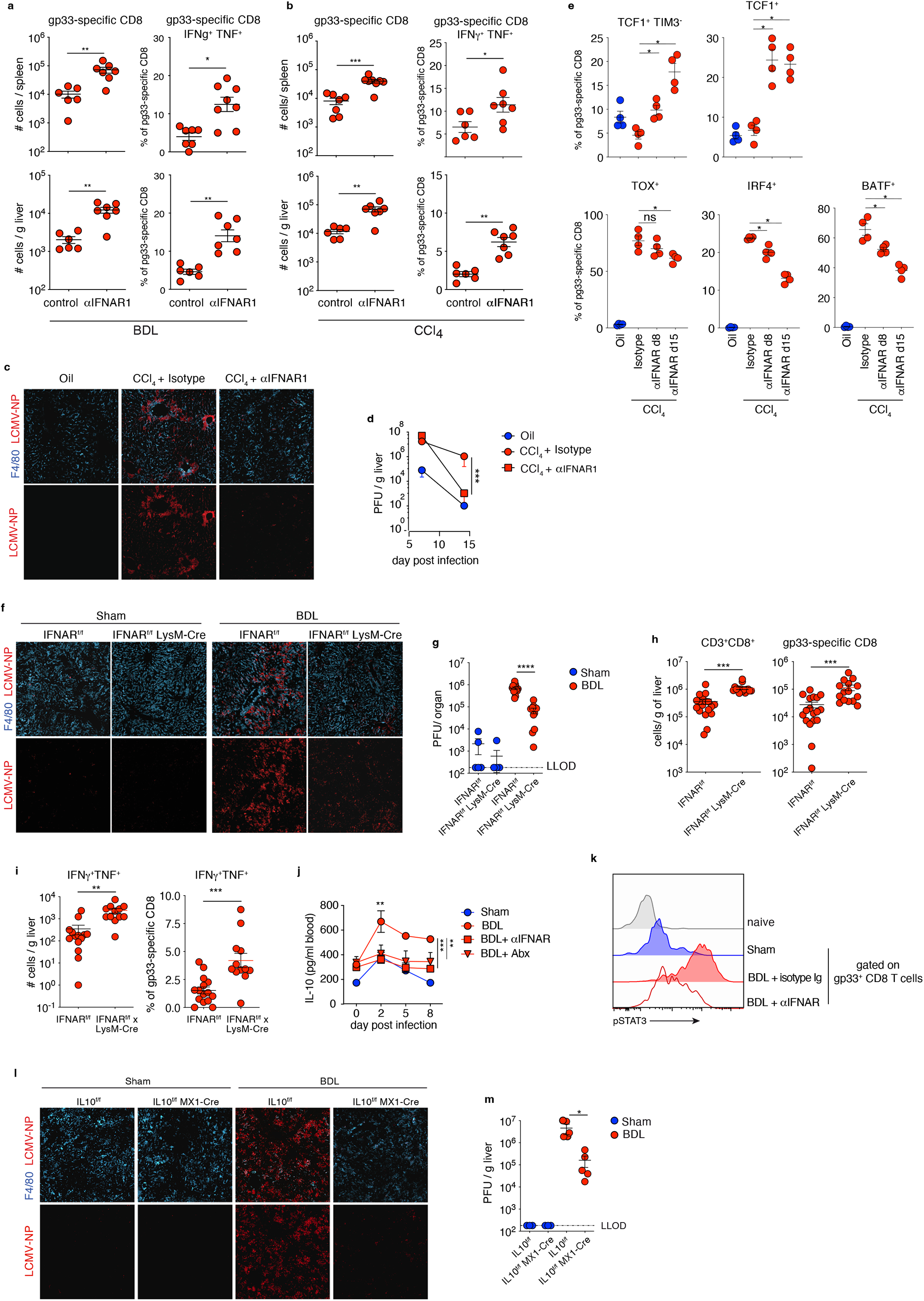
Abrogation of IFNAR-signalling improves functionality of virus-specific CD8 T cell in LCMV-infected BDL-mice. **a**,**b**, numbers of virus-specific and cytokine producing LCMV-gp33-specific CD8 T cells after LCMV infection and treatment with IFNAR-blocking antibody or isotype control Ig from mice with BDL or CCl_4_ induced liver injury (a d8 p.i., b d15 p.i.) (n=5). **c**, immune fluorescence of liver tissue for LCMV nucleoprotein and F4/80^+^ macrophages in CCl_4-_treated or control mice on d15 p.i. (n=5). **d**, quantification of LCMV titers (n=5). **e**, percent of transcription factor expressing LCMV-gp33-specific CD8 T cells d15 p.i. (n=5). **f**-**i**, liver immune fluorescence for LCMV nucleoprotein and F4/80^+^ macrophages in IFNAR^flox/flox^ and IFNAR^flox/flox^xLysm-Cre mice (**f**), LCMV titer quantification (**g**) and number of virus-specific CD8 T cells (**h**) and cytokine-producing T cells (**i**) (n=4). **j**, IL-10 levels in blood of BDL-mice treated with anti-IFNAR antibodies (n=3). **k**, phosphorylated STAT3 in gp33-specific CD8 T cells at d8 p.i. from mice d8 p.i. after IFNAR-blockade. **l**,**m**, liver immune fluorescence for LCMV nucleoprotein and F4/80^+^ macrophages in IL10^f/f^ x Mx1-Cre mice or Cre^negative^ littermates at d8 p.i. of BDL- or sham-operated mice and quantification of LCMV titers. **a-m**, data from three independent experiments. **a**,**b**,**d**,**e**,**g**,**h**,**I**,**m** statistics were assessed by unpaired t-test, **j**, one-way ANOVA with Dunnett’s multiple comparisons test; *P≤0.05, **P≤0.01, ***P≤0.001, ****P≤0.0001.

**Figure 6.**
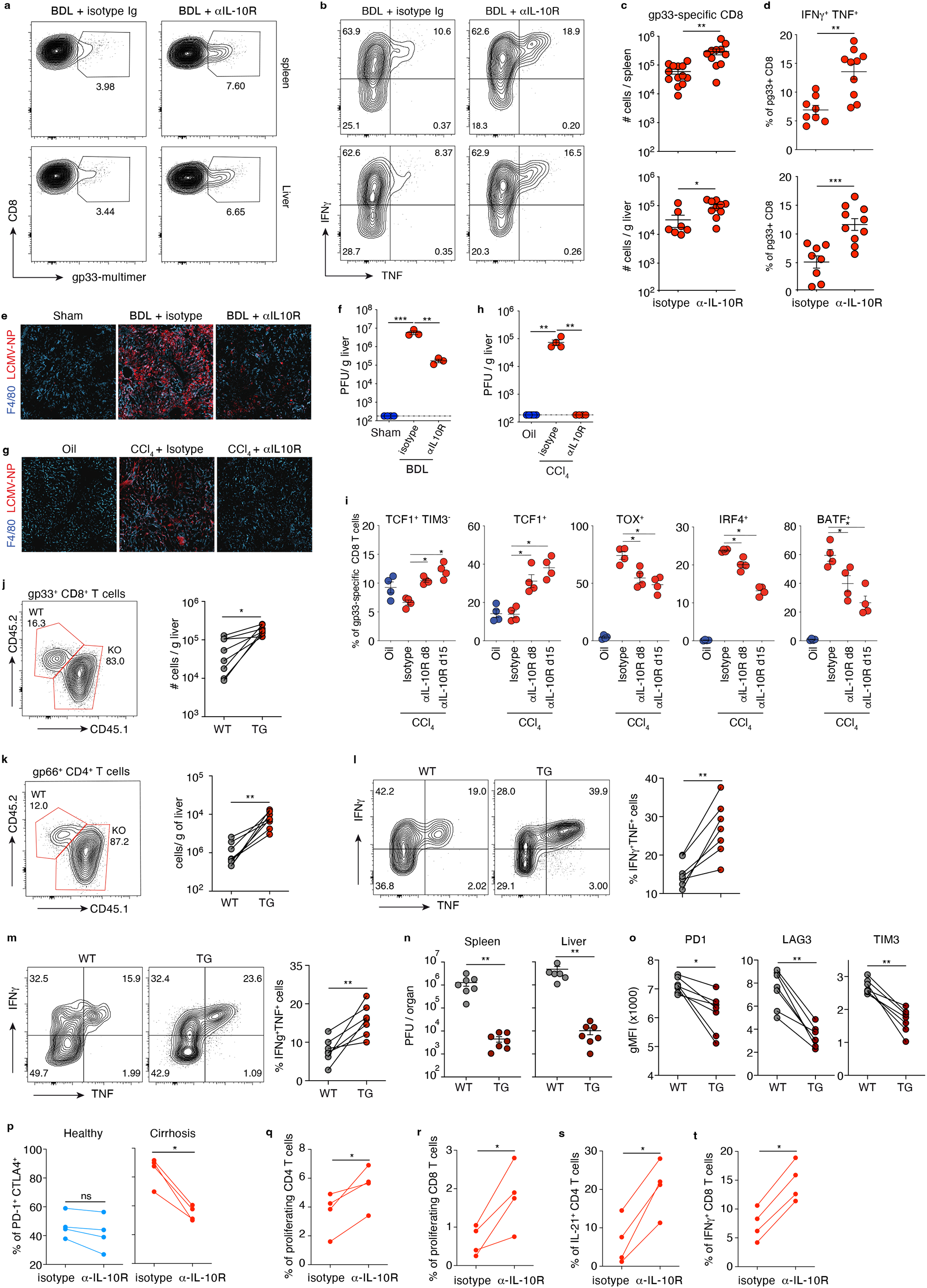
Interference with IL-10 signalling rescues antiviral T cells responses during liver injury. **a-d**, frequencies of LCMV gp33-specific CD8 T cells and cytokine production in BDL mice at d8 p.i. and after anti-IL-10Rα antibody treatment and quantification. **b**,**d**, IFNγ and TNFα CD8^+^ T producing liver CD8^+^ T on day 8 post infection. **e**-**h**, liver immune fluorescence LCMV nucleoprotein and F4/80^+^ macrophages in BDL-mice after anti-IL-10Rα antibody treatment of BDL-mice or CCl_4_ treated mice and quantification of LCMV titers in liver. **i**, transcription factor expression in gp33-specific CD8 T cells. **j-o**, sub-lethally irradiated wildtype B6 mice (CD45.1) were grafted with 1:1 mixture of naive wildtype (WT, CD45.2^+^) and CD4-dominant negative (DN) IL-10Rα transgenic (TG, CD45.1/2^+^) T cells. 4 weeks later mice underwent either sham or BDL operation and infected 9 days later with LCMV. **j**,**k** abundance of gp33-specific CD8 T cells (**j**) or gp66-specific CD4 T cells (**k**) and expression of IFNγ and TNF in CD8 T (**l**) or CD4 T cells (**m**) from spleen at d8 p.i. **n**, LCMV titer in liver and spleen at d8 p.i. of BDL-mice grafted with naive WT or TG T cells. **o**, co-inhibitory molecule expression in LCMV gp33-specific CD8 T cells at d8 p.i. **p**, co-expression of PD1 and CTLA4 by T cells from healthy controls or cirrhotic patients after *ex vivo* anti-CD3 and anti-CD28 stimulation in presence of IL10Rα blocking (red) or isotype control (grey) antibodies. **q-t**, frequencies of proliferating (**q**,**r)** and cytokine producing (**s**,**t)** HBs-specific CD4 and CD8 T cells from healthy controls (n=4) and cirrhotic patients (n=4) after vaccination against HBsAg and *ex vivo* stimulation with a peptide pool covering HBsAg in presence of IL10Rα blocking antibodies or isotype control. **a-o**, data from three independent experiments. **c**,**d** data are pooled from three independent experiments. **p-t** data are derived from two independent experiments. **c**,**d**,**f**,**h**, statistics were assessed by unpaired t-test, **I** by Kruskal-Wallis tests with Dunn’s multiple comparisons tests, **j-t** comparison between groups was calculated using a two-tailed paired student’s t-test, ns not significant, *P≤0.05, **P≤0.01, *P≤0.05, **P≤0.01, ***P≤0.001.

We therefore characterized IFNAR-signaling in myeloid immune cells, which we recently identified as key cell population curtailing anti-bacterial innate immunity during liver injury^18^, and created mice, which lack IFNAR expression in monocytes, neutrophils and macrophages (LysM-Cre *x* IFNAR^fl/fl^ mice). Strikingly, LysM-Cre *x* IFNAR^fl/fl^ mice with liver injury had improved control of LCMV infection and achieved a 55-fold reduction in viral titers compared to IFNAR^fl/fl^ littermates (**Fig. 5f,g**). In the absence of IFNAR-signaling in myeloid cells, we detected higher numbers of LCVM-specific CD8 T cells, which produced IFNγ and TNF after specific re-stimulation *ex vivo* (**Fig. 5h,i Extended Data Fig. 6e,f**). Of note, lack of IFNAR-signaling in myeloid cells also rescued lymphopenia (**Fig. 5h**). Finally, expression levels of PD1, LAG3 and TIM3 in LCMV-specific CD8 T cells were reduced, when myeloid cells lacked IFNAR (**Extended Data Fig. 5g,h**). This indicated that IFNAR-signaling in myeloid cells was a key event for T cell dysfunction during liver injury and raised the question how IFN I stimulated myeloid cells impaired T cell function. The observation that IFNAR-signaling in myeloid cells induces expression of IL-10 upon intracellular infection^18^ led us to study the contribution of IL-10 to T cell dysfunction during liver injury.

We found high IL-10 blood-levels in mice with liver injury that further increased after LCMV infection, but were absent after anti-IFNAR or antibiotics treatment (**Fig. 5j**). Higher IL-10 levels were associated with higher levels of phosphorylated STAT3 (**Fig. 5k**), the canonical downstream signaling molecule of the IL-10 pathway, which was reduced by anti-IFNAR treatment (**Fig. 5j,k**). Furthermore, GSEA showed an enrichment of IL10R-stimulated genes in CD8 T cells in BDL-mice (**Extended Data Fig. 7a**), overall pointing towards a role for IL-10 in impacting on the phenotype and function of T cells in the context of liver injury. We therefore examined the link between IFNAR signaling in myeloid cells, elevated IL-10 expression, and T cell dysfunction during liver injury, and generated MX1-Cre *x* IL-10^fl/fl^ mice, in which the *Il10* gene was excised in cells expressing the IFN-stimulated gene MX1^42^. In MX1-Cre *x* IL-10^fl/fl^ mice with liver injury, LCMV-specific CD8 and CD4 T cells were increased in numbers (**Extended Data Fig. 7b-e**) and showed higher levels of cytokine expression compared to control mice (**Extended Data Fig. 7f-h**). Notably, abrogation of IFNAR-mediated IL-10 expression resulted in improved T cell immunity and reduction of LCMV titers (**Fig. 5l,m**). Overall, these data indicated that IFNAR-induced IL-10 expression in myeloid cells contributed to T cell dysfunction and failure to control viral infection during liver injury.

### IL-10 signaling acts directly on antigen-specific T cells and can be targeted to reinvigorate anti-viral immunity during liver injury

To further investigate the role of IL-10 in modulating antiviral T cell immune surveillance during liver injury, we used anti-IL-10Rα antibodies to abrogate IL-10 signaling (**Extended Data Fig. 8a**). Blockade of IL-10Rα-signaling during LCMV-infection in mice with liver injury led to an increase in the numbers of LCMV-specific CD8 T cells (**Fig. 6a,c, Extended Data Fig. 8b,d**) that co-expressed IFNγ and TNF (**Fig. 6b,d, Extended Data Fig. 8c,e**). Furthermore, after anti-IL-10Rα treatment expression levels of PD-1, LAG-3 and TIM3 on LCMV-specific CD8 T cells decreased (**Extended Data Fig. 8f,g**). Consistent with this quantitative and qualitative improvement of T cell immunity, IL-10Rα blockade in mice with liver injury caused a 100-fold reduction of LCMV titers on day 8 after infection or even clearance of infection (**Fig. 6e-h**). Moreover, treatment with anti-IL10Ra antibodies led to increased frequencies of TCF1-expressing T cells (**Fig. 6i, Extended Data Fig. 8h,i**) and reduced expression levels of TOX, IRF4 and BATF (**Fig. 6i, Extended Data Fig. 8i**), similar to what we observed after IFNAR-signaling blockade. Taken together, these data revealed that blocking IL-10Rα-signaling restored antiviral T cell immune surveillance to control LCMV infection.

To determine whether IL-10 acted directly on T cells or whether its inhibitory effect depended on other cell populations, we analyzed IL-10Rα expression on different immune cell populations. While antigen-experienced CD44^+^CD8 and CD44^+^CD4 LCMV-specific T cells expressed IL-10Rα protein at higher levels in mice with liver injury (**Extended Data Fig. 9a-c**), no differences in IL-10Rα expression were observed for myeloid cells or CD44^neg^ T cells (**Supplemental Data 1. a-d**). Importantly, in cirrhotic patients we also detected elevated levels of IL-10Rα on antigen-experienced CD45RA^neg^ T cells compared to T cells from healthy individuals (**Extended Data Fig. 9d**). Together, these data suggested that T cells might have a higher sensitivity to IL-10 signaling if liver injury was present, and led us to investigate a possible link between tonic IFNAR-signaling and elevated IL-10Rα expression. For this, we treated murine naive T cells with IFN I for 48hrs prior to T cell receptor stimulation *in vitro* or induced IFN I production *in vivo* by the administration of polyI:C for 1-3 days. Pre-exposure of T cells to IFN I significantly *in vitro* or *in vivo* enhanced their responsiveness to the inhibitory functions of IL-10 by upregulating IL-10Rα in a time and dose-dependent fashion (**Extended Data Fig. 9e**,**f**).

To investigate whether IFN I-induced IL-10 impaired immune surveillance by acting directly on T cells, we co-transferred equal numbers of wild type (WT) or CD4–DN IL-10Rα transgenic naïve T cells (TG), which overexpressed a dominant negative (DN) IL-10Rα chain and therefore had impaired IL-10 signaling^43^, into irradiated wild type mice. Co-transfer of WT and TG cells ensured that T cells were exposed to the same microenvironment. After induction of liver injury in chimeric mice (CD45.1^+^) successfully reconstituted with WT (CD45.2^+^) and TG (CD45.1/2^+^) T cells (**Supplemental Data 1e**,**f**), mice were infected with LCMV and T cells were analyzed. Strikingly, in LCMV-infected mice with liver injury total numbers of transgenic (TG) CD8 T cells were 10-fold and 100-fold higher in the liver, compared to WT cells (**Extended Data Fig. 9g**). More importantly, CD4 and CD8 T cells with impaired IL-10Rα-signaling were much more prevalent among LCMV-specific T cells in the liver (**Fig. 6j,k**) and had superior effector cytokine production compared to WT cells (**Fig. 6l,m**). Consistently, after transfer of T cells with impaired IL-10Rα-signaling, LCMV-infected mice with liver injury had 100-fold lower viral titers in liver and spleen than counterparts that received WT T cells (**Fig. 6n**). Thus, T cells with low responsiveness to IL-10 had high proliferation capacity and effector cytokine production during liver injury. Of note, while LCMV-specific T cells with impaired IL-10Rα-signaling (TG) had lower expression levels of PD-1, LAG-3 or TIM3 (**Fig. 6o, Extended Data Fig. 9h**), there were no differences in expression of IRF4, TOX or BATF (**Extended Data Fig. 9i**) suggesting that IFNAR/IL-10-induced T cell dysfunction during liver injury is uncoupled from the above-mentioned exhaustion-mediating transcription factors. Collectively, these results suggested that increased IL-10 presence during liver injury acted directly on T cells that were highly sensitive to IL-10-signaling resulting in T cell dysfunction and impaired viral clearance.

Having identified IL-10 as cause of T cell dysfunction in preclinical models of liver disease, we wondered whether interference with IL-10Rα-signaling would also restore T cell dysfunction in patients with liver cirrhosis. Antibody-mediated blockade of the IL-10Rα in αCD3/CD28 stimulated T cells from cirrhotic patients led to reduced expression of co-inhibitory receptors like PD-1 and CTLA-4 (**Fig. 6p, Extended Data Fig. 9j**). More importantly, antibody-mediated blockade of IL-10Rα-signaling triggered proliferation of HBs-specific T cells from Hepatitis B-vaccinated cirrhotic patients (**Fig. 6q,r, Extended Data Fig. 9k**) and allowed CD4 and CD8 T cells to produce more IL-21 and IFNγ, respectively (**Fig. 6s,t Extended Data Fig. 9 l**). Collectively, these data provide evidence that IL-10Rα-signaling regulates proliferation and effector functions of virus-specific CD4 and CD8 T cells in patients with cirrhosis.

## Discussion

Cirrhosis-associated immune dysfunction is linked to occurrence of infections that can cause loss-of-function of remaining liver tissue and thereby trigger life-threatening liver failure^44-46^, against which no specific therapeutic intervention exists. While intractable bacterial infections pose the most prominent threat to cirrhotic patients^44^, viral infections also cause liver failure and difficult-to-treat infections in these patients^46-48^. Here, we identify in preclinical models of liver injury and in cirrhotic patients the mechanisms, through which liver injury induces suppression of systemic anti-viral T cell immunity (LIST) by an IFN I/IL10 signaling axis and leads to persistence of infection with LCMV strains readily eliminated in the absence of liver injury.

The outcome of immunity to viral infection is determined by virus-intrinsic properties and by host factors influencing anti-viral immunity. Such virus-intrinsic properties and their contribution to development of persistent viral infection have been extensively studied, such as lymphocytic choriomeningitis virus clone 13 infection overcoming anti-viral immunity by inducing T cell exhaustion^15,16,29^, human immunodeficiency virus achieving persistence through viral integration into the host genome of T cells^49,50^, or hepatitis B virus establishing a robust viral latency form in hepatocytes and exploiting the liver’
ss tolerogenic properties^51-55^. On the other side, host factors such as age, inflammation and comorbidities, amongst which anti-viral immune responses develop, also shape the outcome of infection^32,56,57^. In COVID-19 patients with cardiovascular, lung or metabolic diseases overshooting immune responses causing organ pathology are more frequently observed^58,59^. In contrast, in patients with liver cirrhosis attenuated and dysfunctional immune responses are observed^44,60-64^, which points towards a particular role of liver injury in the decreased immunity.

The liver serves as a firewall that eliminates intestinal microbiota that gained access to portal venous blood^65,66^. In liver cirrhosis patients, intestinal dysbiosis in combination with increased microbial translocation is believed to cause systemic inflammation^67,68^. We found that enhanced translocation of intestinal microbiota after liver injury caused tonic IFN-signaling in liver macrophages and that abrogation of IFN-signaling in these cells prevented loss of systemic T cell immunity. However, we did not find evidence for intestinal dysbiosis, and reconstitution of germfree mice with gut microbiota healthy mice or mice with liver injury led to similar levels of IFN I and similar loss of systemic T cell immunity. This suggests that bacterial translocation rather than intestinal dysbiosis was responsible for loss of systemic T cell immunity during liver injury, and raised the question how IFN-signaling in myeloid cells could regulate T cell function.

Genome-wide transcriptional and high-parametric protein profiling of LCMV gp33-specific T cells from injured liver after LCMV infection revealed gene signatures for TGFβ-signaling, IFN-signaling, TNF-signaling and STAT3-signaling among others. Of note, we also detected gene signatures characteristic of exhausted T cells found in cancer or chronic infection. However, loss of T cell effector function during liver injury preceded the upregulation of T cell exhaustion-associated transcription factors like TOX, BATF and IRF4, corroborating our assumption that loss of virus-specific T cell immunity during liver injury is not a cell-intrinsic process but is driven by myeloid cells in a paracrine fashion. Strikingly, blockade of TGFβ-signaling *in vivo* led to severe hepatic immune pathology but no reduction in LCMV titers identifying a non-redundant role of TGFβ in organ-protection but not in antiviral immunity. The IL10-gene signature and enhanced STAT3-signaling prompted us to explore a role of IL-10 predominantly produced by monocytes/macrophages^69^. Importantly, after LCMV-infection of mice with liver injury we detected increased IL-10 expression in monocytes/macrophages that depended on IFN I-signaling, which is unexpected because IFN I is involved in self-perpetuating inflammation^70^.

In contrast to the immune pathology-promoting effect of blocking TGFβ, inhibition of IL-10 by antibodies or T cell-specific ablation of IL-10 receptor signaling rescued effector function of LCMV-specific CD8 T cells in mice with liver injury and led to control of viral infection, nota bene in the absence of immune pathology. Likewise, elimination of intestinal microbiota and reducing gut microbial translocation to the liver as well as blockade of IFN-signaling in myeloid cells all prevented expression of IL-10 and rescued antiviral T cell function. Our results also point towards a role of liver injury in chronic viral infection of the liver, where virus-specific T cells fail to eliminate infected hepatocytes^55^. Translating these results from preclinical models of liver injury to patients with liver cirrhosis, we found that inhibition of IL-10 receptor signalling rescued vaccination-induced antigen-specific T cells from their dysfunction.

Our work therefore provides a mechanistic understanding of the loss of systemic T cell immunity during liver injury, discriminates organ-protective TGFβ-signalling from T cell suppressing IL-10 signalling and identifies IFN I and IL-10 as molecular targets for immune interventions that reconstitute T cell immunity without immune pathology.

## Methods

### Data availability

All data from this study are provided in the source data.

RNA-seq expression data are available at NCBI GEO under the accession number: *GSE158261*. Primary data from flow cytometry or immunohistochemistry are available upon reasonable request.

### Patients and blood samples

Blood and serum samples as well as questionnaire-based assessment of donor characteristics and disease stage from cirrhosis patients and healthy volunteers before vaccination with TWINRIX (HAV and HBs Antigen of the HBV) were collected at the University Hospital Bonn and general practices in Bonn (healthy individuals; n = 16 and cirrhotic patients n = 16). All enrolled subjects were never exposed to HBV or been vaccinated against HBV or HAV, proven by the absence of anti-HBs antibodies by serological testing prior to vaccination and patient information. Informed consent was obtained from all patients and healthy individuals enrolled in the trial in accordance with the Declaration of Helsinki protocol. The study was approved by and performed according to the guidelines of the local ethics committees of the University of Bonn (310/16). All enrolled subjects received 4 doses of the vaccine on d1, d7, d21 and after 6 months. Sample collection was performed approximately 4-5 weeks after the last injection. PBMCs were isolated by Ficoll-Hypaque (PromoCell, Germany) density gradient centrifugation and stored at −80 °C until further use. Serum was separated by centrifugation for 10 min and supernatant was stored at −80 °C. Detailed cirrhosis and healthy donor characteristics and the study design are provided in Supplementary Table I and Extended Data Fig. 1.

### HBS-specific T cell activation

43 (80-90% purity) 15-mer synthetic peptides overlapping by ten amino acids overlap (Xaia Custom Peptides, Sweden) covering the sequence of the HBV surface antigen (HBs Antigen) protein according Galibert sequence^71^ (Supplementary Table 2). Peptides were pooled and used for detection of HBs-specific T cells and their function. After isolation from peripheral blood, untouched CD3 T cells were labelled with cell trace violet (CTV) (0.5 μM; Thermo Fisher Scientific) and cultured in round-bottom 96-well plate at 5 × 10^4^/well, in the presence of autologous irradiated (4,000 rads) PBMC (10^5^/well), HBs-peptides and recombinant human IL-2 (50 U/ml; PeproTech). After 5 days of incubation, proliferation and cytokine production in HBS-specific T cells were assessed by flow cytometry.

### Anti-RBD (S) SARS-CoV-2 antibodies assay

Serum samples were analyzed for anti-S-RBD IgG titers using the SARS-CoV-2 Plate 7 V-PLEX Serology Kit from MSD (Meso Scale Diagnostics, LLC). Antibody concentrations were quantified via the MESO SECTOR S 600. Raw data was analyzed with the MSD Discovery Workbench tool (V 4.0.13), that quantifies anti-RBD IgG. All assays were performed by trained laboratory technicians according to the manufacturer’s standard procedures.

### In vitro activation and proliferation assay of human T cells

PBMCs were labelled with CTV (0.5 μM; Thermo Fisher Scientific) and cultured in RPMI supplemented with 10% FCS at 1 × 10^6^ cells/well in 96-well plates (Corning) coated with anti-CD3 (clone HIT3a, BioLegend) and anti-CD28 (clone L293, BD Biosciences) (each at 3 μl/ml) and IL-2 (10 IU/ml; PeproTech). T-cell proliferation and intracellular cytokine expression as well mitochondrial assays were assessed by flow cytometry.

### In vitro IL-10Rα blockade on human T cells

PBMCs were cultured in RPMI supplemented with 10% FCS at 1 × 10^6^ cells/well in 96-well plates (Corning) coated with anti-CD3 (clone HIT3a, BioLegend) and anti-CD28 (clone L293, BD Biosciences) (each at 3 μl/ml) and IL-2 (10 IU/ml; PeproTech). 5μg/ml of the blocking anti IL-10Rα antibody (3F9; BioLegend) was added 12 hours later.

### ELISA for detection of cytokine expression

Human IFNγ, TNF and IL-2 in the supernatant of *in vitro* activated cells and murine IL-10 in the serum were determined using ELISA MAX Deluxe Set (BioLegend) according to manufacturer instructions.

### Quantification of mitochondrial membrane potential, mitochondrial mass and mitochondrial superoxide production

For mitochondrial studies in human T cells were determined after anti-CD3 stimulation. Cells were incubated with 50 nM MitoTracker Green (MTG) and/or 25 nM MitoTracker DeepRed (MTDR) for 30 min at 37C prior to cell surface staining. Mitochondrial superoxide levels in human T cells were determined, after cell surface staining, by incubation (15 min at 37 °C) of overnight anti-CD3 stimulated and unstimulated cells in the presence of MitoSOX Red (5μM; Molecular Probes).

### Mice

6-9 week old mice C57BL/6J (B6) were purchased from Janvier (Le Genest-Saint-Isle, France. LysM^Cre^-IFNAR^fl/fl^ (previously described^18^), OT-I, OT-II, CD45.1-B6 (B6.SJL-Ptprca Pepcb/BoyJ), and P14 mice were originally purchased from The Jackson Laboratory and maintained in the House of Experimental Therapy, University Clinic Bonn. Mx1^Cre^-Il10^fl/fl^ x Mx1^Cre^ ((Il10^tm1Roer^) x (C.Cg-Tg(Mx1-Cre)1Cgn/J)) were kindly provided by Axel Roers. IL10Ra^DN^ mice^43^ were kindly provided by Samuel Huber. All mice were maintained under specific pathogen-free conditions specific pathogen-free (SPF) conditions and were handled according to the guidelines of the to institutional animal guidelines of the animal facilities of the University of Bonn. Experimental procedures were approved by the Animal Ethics Committee of the state of North Rhine-Westphalia, Germany. For antibiotic treatment, mice were given a combination of vancomycin (1 g/l), ampicillin (1 g/l), kanamycin (1 g/l), and metronidazole (1 g/l) in their drinking water^72^. All antibiotics were obtained from Sigma Aldrich. Colonization of Germ Free (GF) mice with caecal microbiota of SPF mice was perform as previously described ^73^. Briefly GF mice received 3 times a suspension of the cecum content of sham or BDL operated wildtype per oral gavage on 3 consecutive days and were BDL operated one week after the last transfer.

### Murine liver injury models

Liver injury was induced in male, 8-9 week old mice via bile duct ligation (BDL) or treatment with carbon tetrachloride (CCl4) following established protocols^*74,75*^. In brief, to induce BDL, the animals were treated with painkillers and anaesthetized before peritoneal cavity was opened along the *linea alba*. Two ligatures were placed around the common bile duct in order to obstruct it, before the incisions in the peritoneum and the skin were then closed and the mice were allowed to recover. During the first 5 days after operation all animals received additional injections of painkillers and liver injury was allowed to develop for 10 days before experiments were performed. Alternatively, mice received 0.5 μl CCl_4_ /g body weight for 12 weeks intraperitoneally. CCl_4_ was dissolved 1:7 in olive oil and was administered in 3-day intervals. After the final injection, mice were allowed to recover for 10 days before further experiments were performed. Control mice were sham-operated (no ligation of the bile duct) or received olive oil (i.p.) injections respectively. During all experiments, animals were monitored closely on a daily basis. To inhibit of IFNAR or IL10Rα signaling *in vivo*, mice were treated intraperitoneally with 250μg/mouse of blocking antibodies targeting TGFBR-II (clone: 1D11.16.8) IFNAR1 (clone: MAR1-5A3) or IL10Rα (clone: 1B1.3A) from BioXcell intravenously. Control animals received injections containing the HPRN and MOPC-21 antibodies respectively.

### LCMV infection

Mice were infected with 2×10^4^ plaque-forming units (PFU) of LCMV strain WE or Armstrong diluted in sterile PBS intravenously.

### Irradiation and adoptive cell transfers

Mice were subjected to sublethal irradiation (6 Gy) one day before transfer. T cells were isolated from the spleens and lymph nodes of donor mice and 4×10^6^ CD3^+^ T cells (1:1 of WT and TG cells) per recipients were transferred intravenously. Prior to any subsequent experiment, recipient mice were allowed to recover for 12 days in order to ensure proper engraftment. For all other infection or vaccination studies, Naïve P14, OT-I and OT-II cells were isolated from the spleens of mice and 2.5×10^5^ cells were transferred one day prior to vaccination.

### Murine in vitro T cell proliferation assay

48-well plates were coated with antibodies directed against murine CD3 (clone: 500A.2, BD Biosciences) and CD28 (clone: 37.51, Biolegend) at a concentration of 0.5μg/ml and 10μg/ml respectively, incubated at 37°C for 2 h and washed. T cells isolated from the spleens of Sham-or BDL operated mice were stained with Cell Trace Violet (Invitrogen) and added at a concentration of 2×10^6^ cells/ml. T-cell proliferation was measured after three days by flow cytometry.

### Flow cytometry and antibodies

Single cell suspensions were acquired on a FACSCanto II or LSRII Fortessa and analyzed with FlowJo (version 10.0.7, Tree star). In order to determine the expression of surface molecules, cells were stained on ice for 20 min. The LIVE/DEAD fixable Near-IR Deal Cell Stain kit (Life Technologies) was used in all staining to detect dead cells, also, an in-house made antibody (clone: 2.4G2) directed against the epitopes shared by Fc-gamma receptors was added. Single cell suspension from spleen or liver were stimulated with 100ng/ml PMA, 200ng/ml in the presence of Brefeldin A and monensin for 3 hours, before collecting and intracellular staining for flow cytometry. To block unspecific staining. Cytokines were stained after fixation with 4% PFA and Permeabilization with 1x Permeabilization buffer (FoxP3/Transcription Factor Staining Buffer Set, eBioScience) and transcription factors with the FoxP3/Transcription Factor Staining Buffer Set according to the manufacturer’s instructions. Antibodies directed against the following targets in mice were purchased from Biolegend, eBioScience or Miltentyi: anti-CD3e (145-2C11), anti-CD4 (GK1.5 or RM4-5), anti-CD8a (53-6.7), anti-CD11b (M1/70), anti-CD11c (N418), anti-CD44 (1M7), anti-CD45.1(A20), anti-CD45.2(104), anti-CD146 (ME-9F1), anti-CD210a (1B1.3a), anti-CTLA-4 (UC10-4B9), anti-Eomes (Dan11mag), anti-F4/80 (BM8), anti-GzmB (GB11), anti-IFNγ (XMG1.2), anti-IL-2 (JES6-5H4), anti-IL-21 (FFA2), anti-LAG3 (C9B7W), anti-PD-1 (EH12.2H7), anti-T-bet (4B10), anti-TCRβ (H57.597), anti-TIM3 (RMT3-23) and anti-TNF (MP6-XT22), p anti-hosphor-SMAD2 (Ser250 (SD207-1)). CD8 and CD4 T cells specific for the LCMV epitope gp33-41 and gp66-77 respectively, were purchased from Immudex (gp33-dextramer) or provided by the NIH Tetramer Core Facility (Emory University). The following antibodies were used to detect protein expression on human cells: anti-CD3 (SK3), anti-CD62L (DREG-56), anti-CD8 (PRA-T8), anti-CD45RA (HI100), anti-CD45 (HI30) from ThermoFisher and anti-CD4 (PRA-T4), anti-PD1 (EH12.2H7), anti-CTLA4 (BNI3), anti-CD3 (OKT3), anti-IFNγ (4S.B3), anti-IL-21 (3A3-N2) anti anti-IL-10Rα (3F9) from BioLegend.

### Immunofluorescence microscopy

Liver tissue samples were fixed in a 0.05 M phosphate buffer containing, 0.1 M L-lysine, 2 mg/ml NaIO_4_, and 10 mg/ml paraformaldehyde (PLP) at pH7.4 overnight. Subsequently, samples were washed in phosphate buffer and dehydrated in 30% Sucrose overnight. Finally, samples were place int Tissue TEK (Sakura Finetek), snap-frozen and stored at -80° C. 20μm sections were acquired on a CM3050S cryostat, rehydrated and stained with antibodies in buffer containing 1% normal mouse serum. Pictures were acquired on an LSM 710 confocal Microscope (Zeiss). A BV421-conjugated antibody directed against F4/80 (BM8) was purchased from Biolegend, for detection of the LCMV nucleoprotein an unconjugated rat-anti-LCMV antibody (VL4) was purchased from BioXCell; expression was detected via an AF647-conjugated goat-anti-rat antibody (Invitrogen).

### RNA extraction, cDNA-synthesis and RT-PCR

Small samples of liver tissue were homogenized in 1ml Quiazol and total RNA was isolated using the RNeasy Lipid Tissue Kit (Quiagen, 74804). Afterwards, 2-5μg of RNA was reverse transcribed into cDNA at 37°C for 2 hours using the High-Capacity cDNA Reverse Transcription Kit (Applied Biosystems, 4368814). cDNA was stored at -20°C and real-time PCR was performed using Taqman primers and probes for *Il10, Ifnb1, Mx1, Hprt* and *Gapdh* in murine samples and *PDCD1, CTLA4, CD244, EOMES, BATF, GAPDH* and *HPRT* in human samples. The relative mRNA expression was calculated with the ΔΔCt-method.

### RNA sequencing and data analysis

Transcriptomic differences in isolated T cells from BDL-treated or sham control mice were determined by QuantSeq 3’mRNA sequencing (Lexogen). FACS sorted cells were lysed in 700μl Trizol and stored until RNA extraction was performed with the RNeasy micro kit (Qiagen). Library production for 3’-mRNA sequencing was performed with up to 160 ng purified RNA according to the manufacturers’ protocol and sequenced on a HiSeq2500 (Illumina) with a sequencing depth of 15 Mio reads per sample (NGS Core Facility, University Hospital, Bonn, Germany). The alignment was performed with STAR (v2.5.3a) against the murine reference genome mm10. Transcripts were quantified with the Partek E/M algorithm and further processed for normalization in R (v3.5.0) with the DEseq2 algorithm (v1.20.0). The data set was further optimized by flooring transcripts with minimal gene counts at least to ≤1 and the exclusion of transcripts with a mean expression ≤10 in every test condition. Differentially expressed genes were identified in the Partek Genomics Suite (v7.18.0402) for T cells in BDL versus sham control cells by a one-way-ANOVA (fold-change |1.5|, FDR-adjusted p-Value ≤0.05). Data visualization and biological interpretation were performed with the Partek Genomics Suite, ClueGo plugin (v2.5.2) for Cytoscape (v3.7.2) and R packages ggplot2 (v3.2.1), EnhancedVolcano (v1.6) and tidyr (v1.0.2). Heatmaps of two groups were created using means across the two groups, with expression being centered around 0 and visualized with Mayday^76^(v2.14).

### Statistical analysis

To determine statistical differences, a two-tailed unpaired or paired Student’s t-test was used when two groups were compared; a repeated-measurements one-way ANOVA was used when three groups were compared. For non-parametric data, the Mann-Whitney test was used when comparing two groups and the Kruskal-Wallis test when comparing three or more groups, respectively. Analysis was done with Prism 8. Statistical significance was set at p < 0.05.

### 16S qPCR for quantification of bacterial DNA

DNA was extracted from samples using MoBio PowerSoil kit (Qiagen). DNA concentration was calculated using a standard curve of known DNA concentrations from E. coli K12. 16S qPCR using primers identifying different regions of the V6 16S gene was performed using Kappa SYBR fast mix. Absolute number of bacteria in the samples was then approximated as DNA amount in a sample/DNA molecule mass of bacteria.

**Supplementary Table 1.**
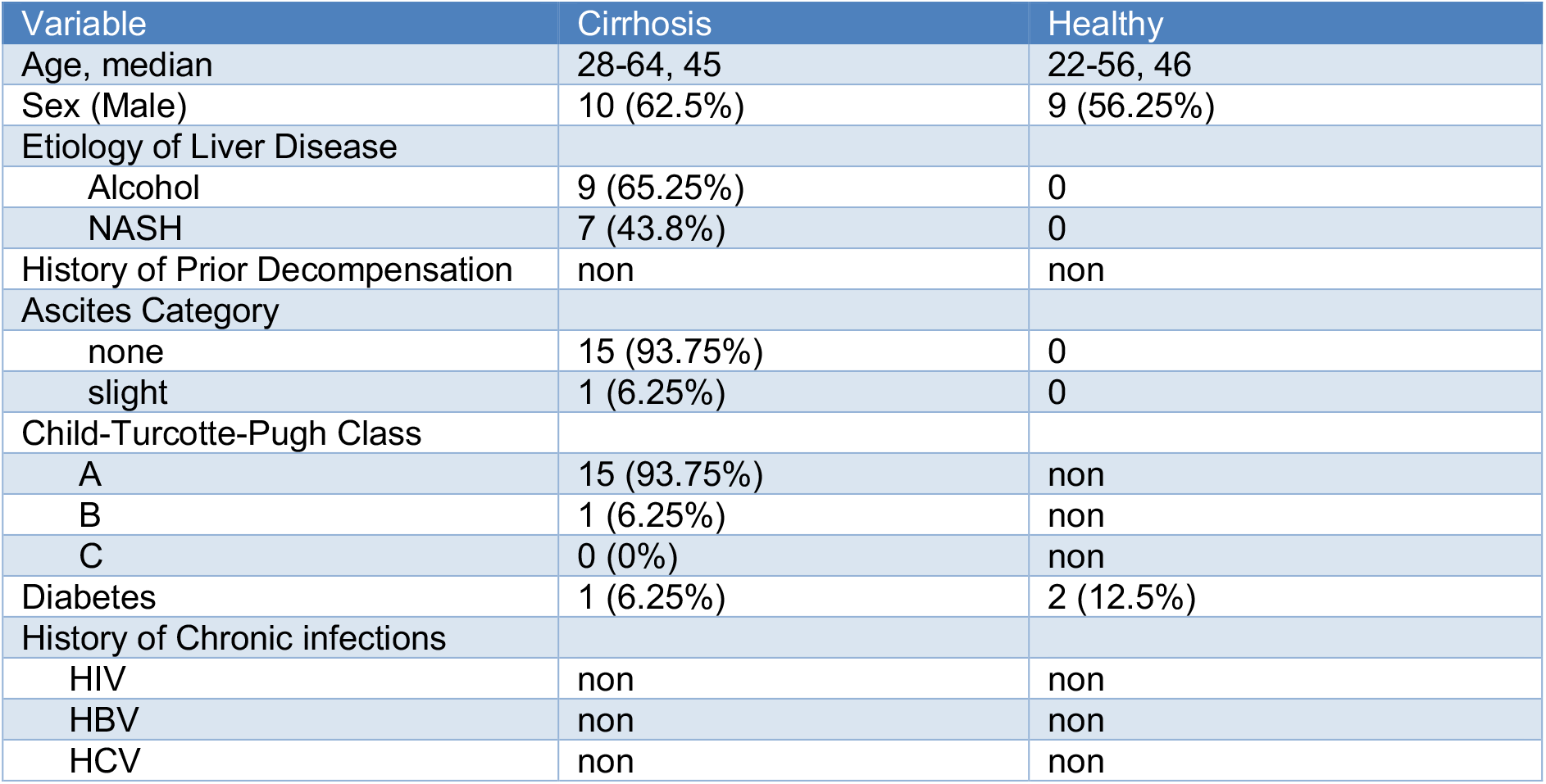

**Supplementary Table 2.**
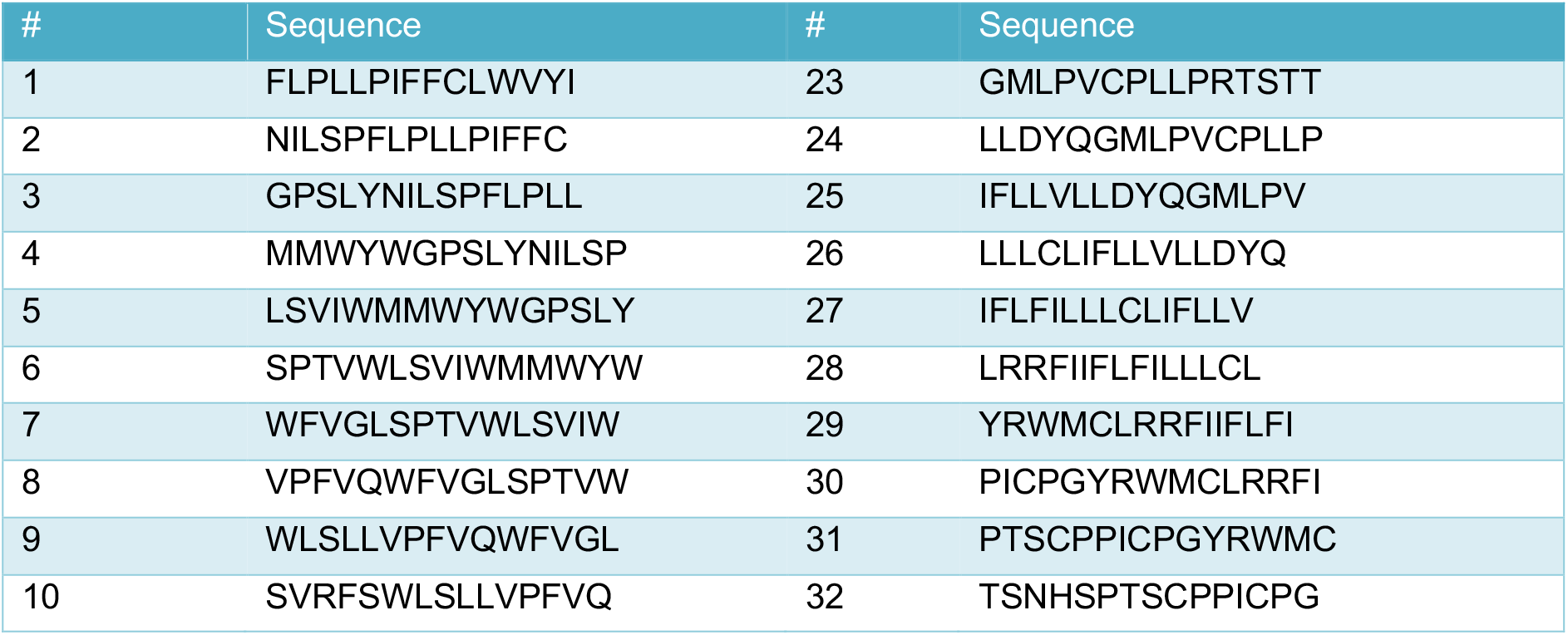
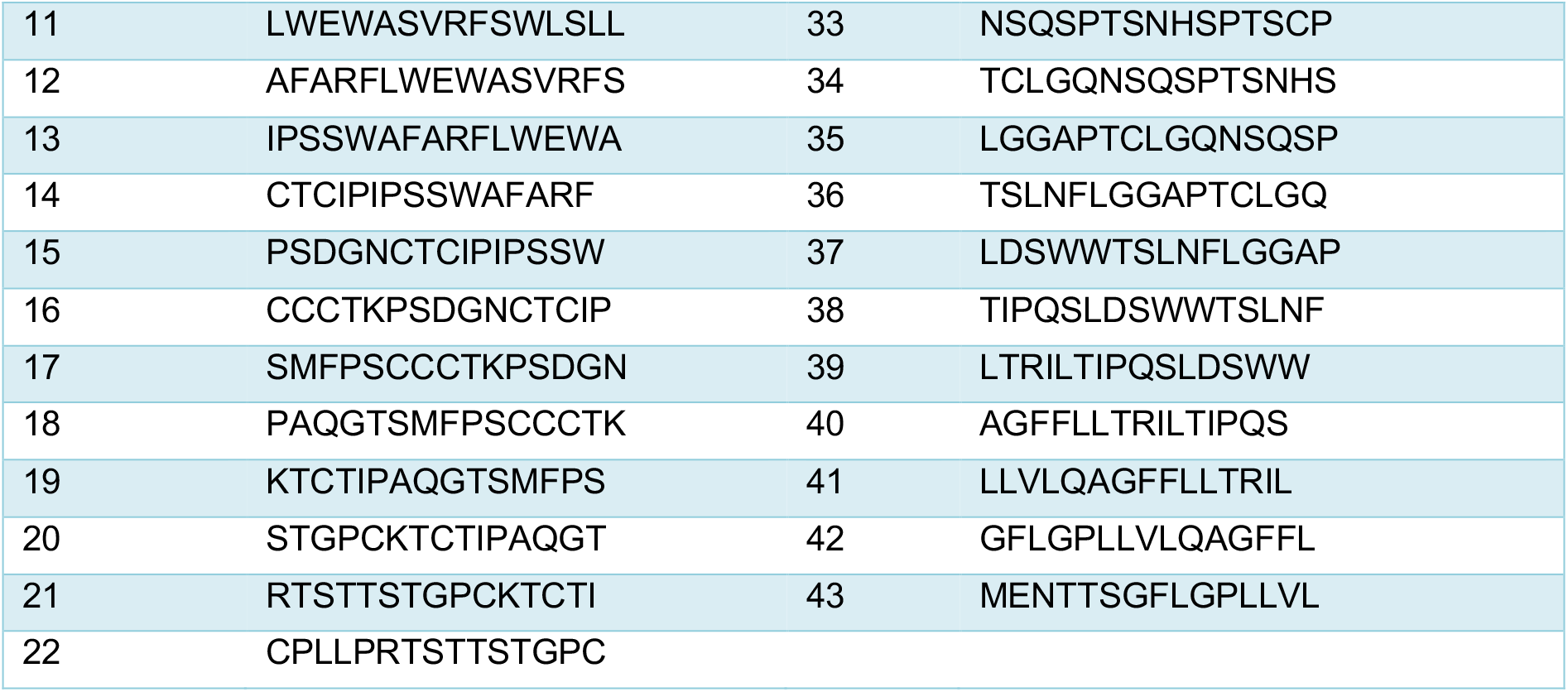

**Supplementary Table 2.**

| Variable | Cirrhosis | Healthy |
| --- | --- | --- |
| Age, average | 35-60, 43 | 36-62, 45 |
| Sex (Male) | 17 (54.8%) | 15 (48.4%) |
| Etiology of Liver Disease |  |  |
| Alcohol | 23 (74,2%) | 0 |
| NASH | 8 (25,8%) | 0 |
| History of Prior Decompensation | non | non |
| Ascites Category |  |  |
| none | 29 (93.5%) | 0 |
| slight | 2 (6.5%) | 0 |
| Child-Turcotte-Pugh Class |  |  |
| A | 27 (87%) | non |
| B | 3 (9.7%) | non |
| C | 1 (3.2%) | non |
| Diabetes | 2 (6.5%) | 3 (9.6%) |
| History of Chronic infections |  |  |
| HIV | non | non |
| HBV | non | non |
| HCV | non | non |

## Acknowledgments

We thank Alisa Ismaili, Rebecca Balduf, Maike Kreutzenbeck and Sven Kröcker for technical assistance. We would like to acknowledge the support by the Central Animal Facility, the Next Generation Sequencing Core Facility, Imaging Core Facility and the Flow Cytometry Core Facility of the Medical Faculty at Bonn University.

## Funding

ZA, PAK, CK, NG, AR, SVS, SH were supported by the Deutsche Forschungsgemeinschaft (DFG). ZA, PAK, NG, KC are funded by the DFG Excellence Cluster (EXC 2151) 390873048. ZA, PAK, JT, CK were supported by SFBTR57. ZA, NG, AR and CK were supported by the SFBTR237. ZA, NG, SVS, CK were supported by the GRK 2168. CK and SH were supported by SFB1192. CPH was supported by the DFG (project number 403193363) and currently receives funding by the Wellcome Trust (WT109965MA, awarded to Paul Klenerman, University of Oxford, United Kingdom). PAK was supported by the European Union’s Horizon 2020 research and innovation programme TherVacB, by the SFB-TRR179 and the German Center for Infection Research, Munich site.

## Notes

The authors declare that they have no conflict of interest

### Competing Interest Statement

The authors have declared no competing interest.

