## Supplemental Figures for "Suppression of systemic T cell immunity to viral infection during liver injury is prevented by inhibition of interferon and IL-10 signaling": supplementary material and figures.pdf

**Extended Data Figures 1 – 9**

**Legends Extended Data Figure 1 – 9**

Extended Data Figure 1

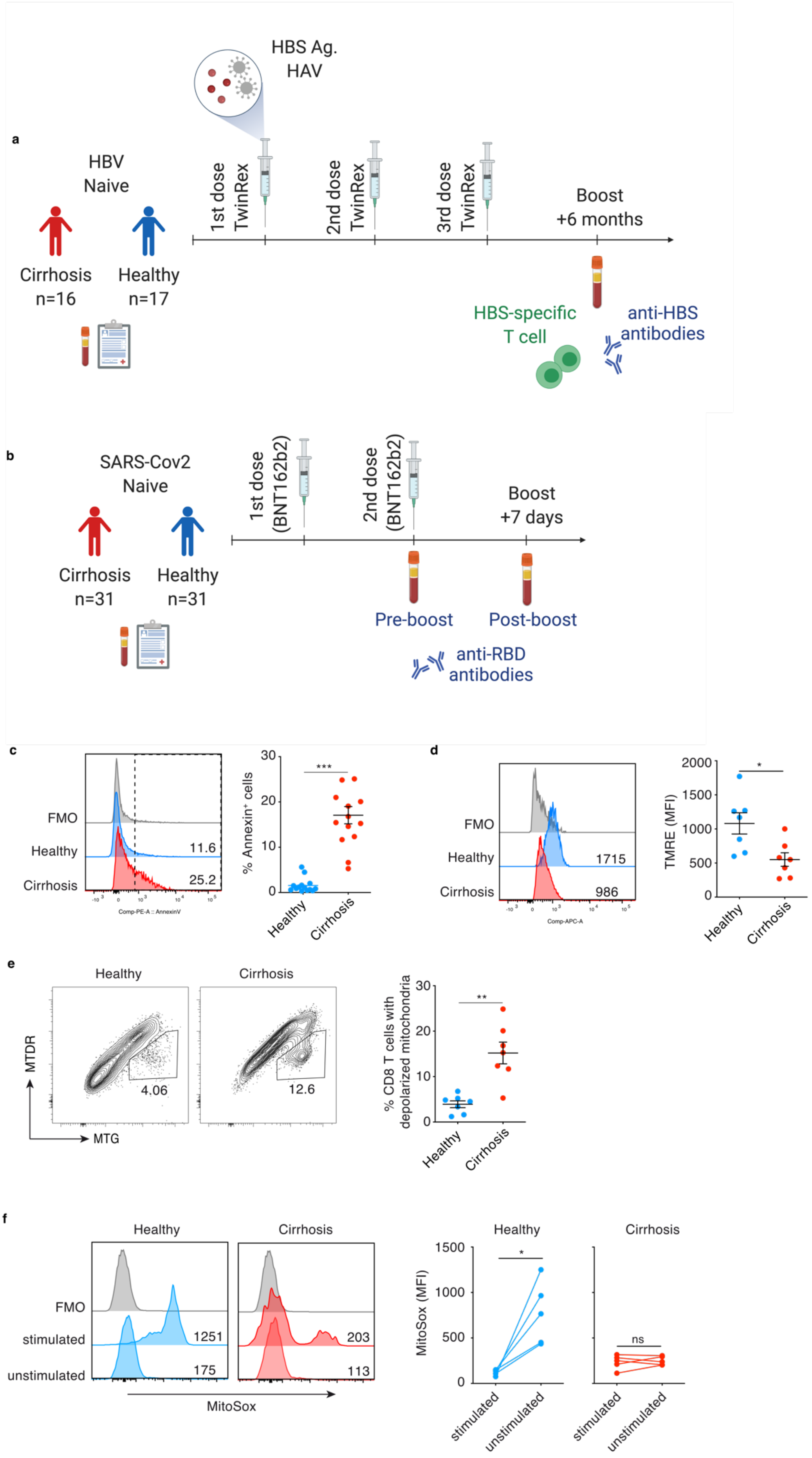

**Extended Data Figure 1 | Design of vaccination study in cirrhotic patients and functional phenotyping of T cells from vaccinated cirrhotic patients**

Schematic of study design (generated by Biorender) for comparison of immune responses to **a**, HBV (TWINRIX) vaccine and **b,** Pfizer-BioNTech mRNA (BNT162b2 - Comirnaty) vaccine in healthy controls and cirrhotic patients. **c-f,** $CD45RA^{+}CD3^{+}$ T cells from healthy subjects or cirrhotic patients and *ex vivo* anti-CD3 and anti-CD28 stimulation evaluated for AnnexinV staining; **c**, staining with mitochondrial dyes TRME; **d**, MTDR; **e**, MTG; **f,** Mitosox. Data are representative of four independent experiments. statistical analysis using a two-tailed paired Student's t-test, ns not significant, *P≤0.05, **P≤0.01, ***P≤0.001.

Extended Data Figure 2

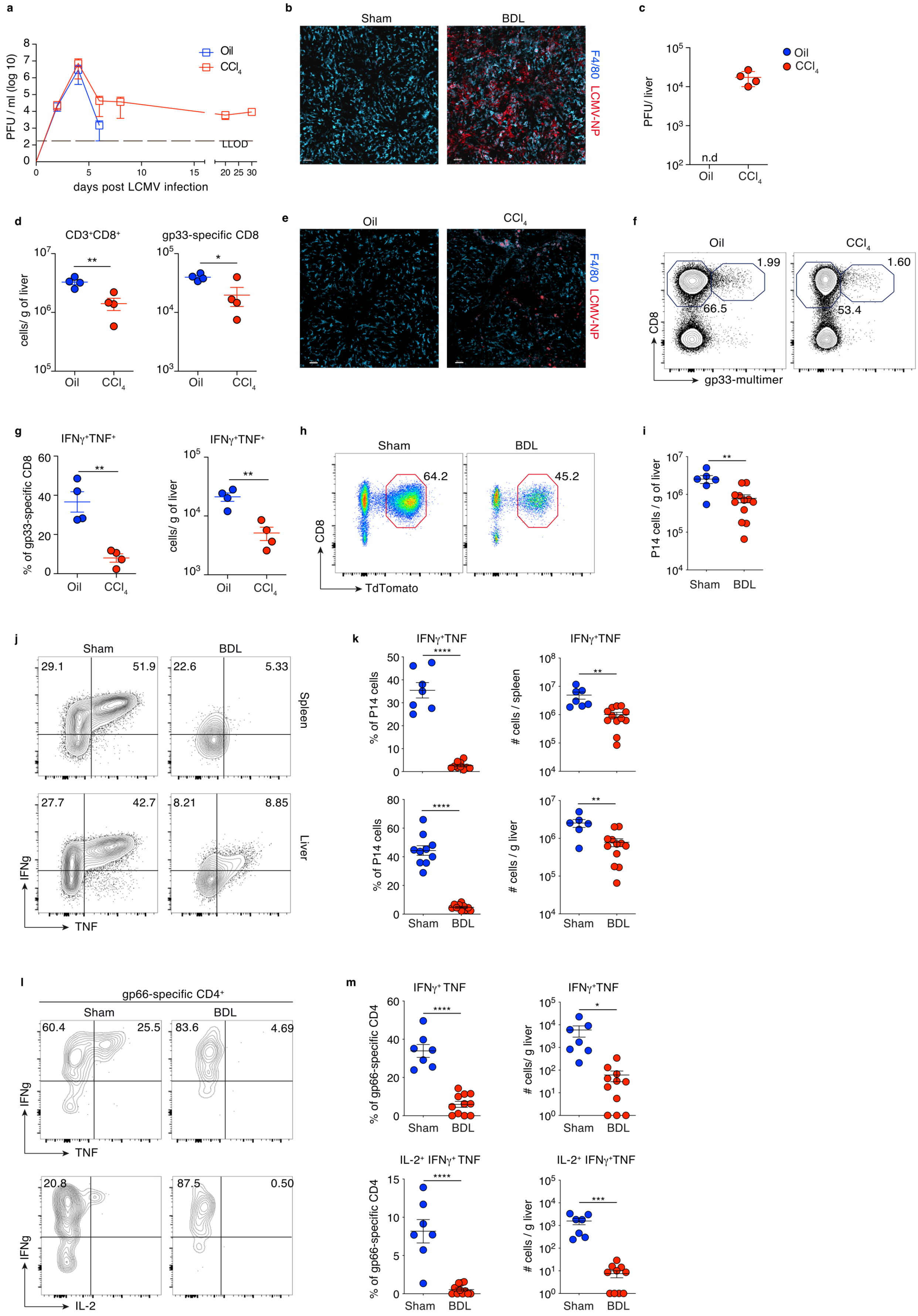

**Extended Data Figure 2 | Functional characterization of anti-viral T cell responses in preclinical models of liver injury in mice**

**a**, LCMV titers (PFU) in blood, liver and spleen of oil- or $CCL_4$-treated mice. **b**,**c**, liver immune fluorescence for LCMV nucleoprotein and F4/80 at d8 p.i. of BDL- or sham-operated mice and quantification of LCMV titers. **d**, numbers of CD8 T cells and LCMV gp33-specific CD8 T from liver of $CCl_4$- or oil-treated mice at d8 p.i. **e**, liver immune fluorescence for LCMV nucleoprotein and F4/80 at d8 p.i. of $CCl_4$- or oil-treated mice at d8 p.i. **f**, frequencies of LCMV gp33-specific CD8 T cells in $CCl_4$- or oil-treated mice at d8 p.i. **g,** percent IFNγ– and TNF-producing LCMV gp33-specific CD8 T cells from liver of $CCl_4$- or oil-treated mice at d8 p.i after *ex vivo* PMA/ Ionomycin stimulation. **h**,**i**, abundance of P14 cells in livers of BDL-mice at d8 p.i. **j,k**, frequencies of IFNγ– and TNF-producing LCMV-specific P14 T cells from liver and spleen of BDL- or sham-operated mice after *ex vivo* PMA/Ionomycin stimulation. **l**,**m**, frequencies and numbers of IFNγ–, TNF- and IL-2 producing LCMV gp66-specific CD4 T cells from liver of BDL- or sham-operated mice after *ex vivo* PMA/Ionomycin stimulation. **a**,**c,d,g,** data from two independent experiments. **i**,**k**,**l,** data pooled from three independent experiments. statistics were assessed by unpaired t-tests, *p≤0.05, **p≤0.01, ***≤0.001, ****≤0.0001.

Extended Data Figure 3

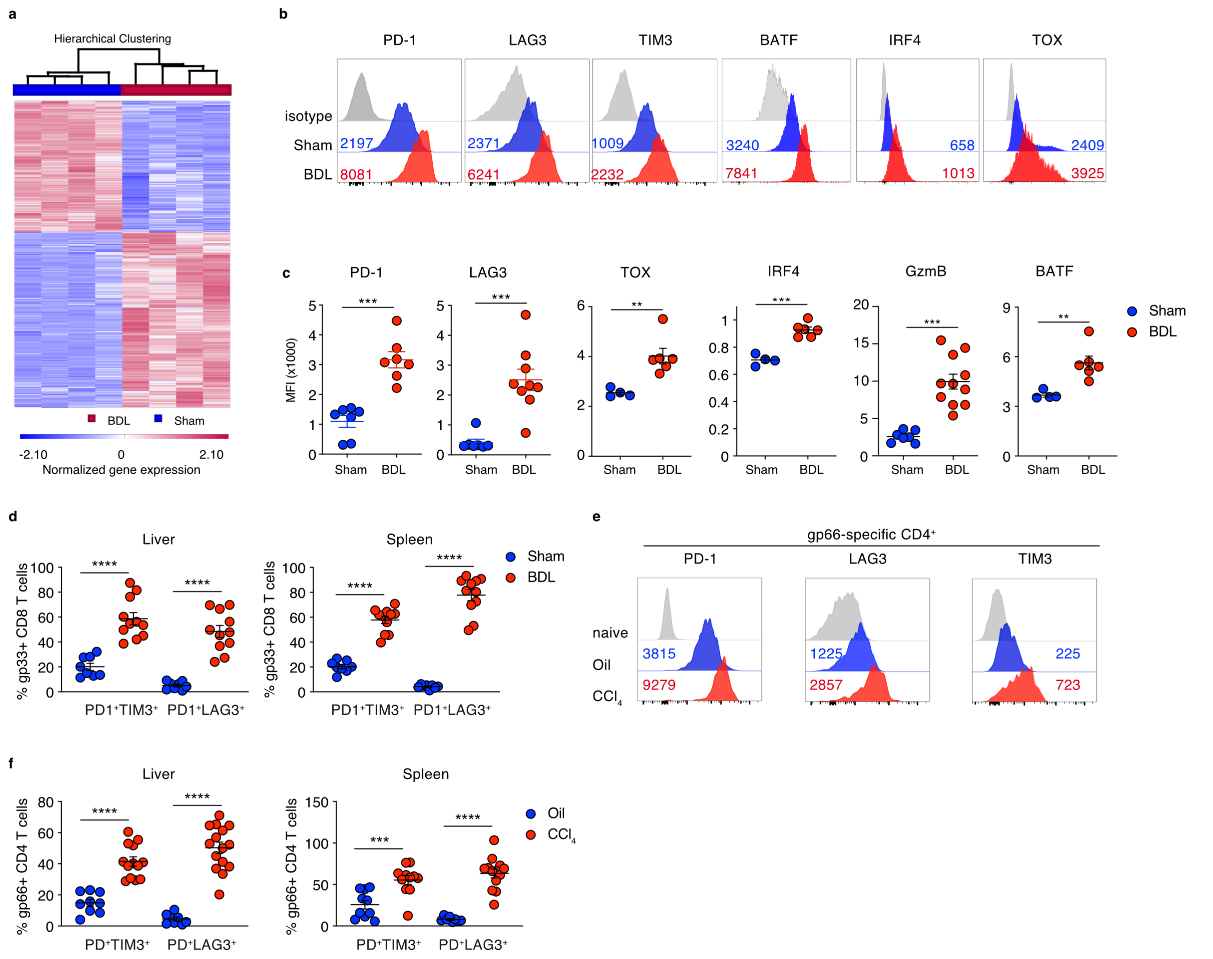


**Extended Data Figure 3 | Transcriptome and molecular profiling of dysfunctional virus-specific CD8 T cells**

**a**, heatmap of 2153 differentially expressed genes (DEGs) in LCMV-specific CD8 T cells BDL- or sham-operated mice; fold-change |1.5|, one-way ANOVA according to Benjamini-Hochberg FDR corrected p-value <0.05. **b**, **c**, histograms and scatter plots showing marker expression by LCMV gp33-specific CD8 T cells from liver of BDL- or sham-operated mice at d8 p.i. **d**, PD-1/TIM3 or PD1/LAG3 co-expression from T cells shown in b. **e**, PD-1, LAG3 and TIM3 expression levels by LCVM gp66-specific CD4 T cells from liver of $CCl_4$- or oil-treated mice d8 p.i. **f**, PD-1/TIM3 or PD1/LAG3 co-expression from cells in e. **b-f**, data from three independent experiments. **c,d,f**, statistics were assessed by unpaired t-tests, **P≤0.01, ***P≤0.001, ****P≤0.001.

Extended Data Figure 4

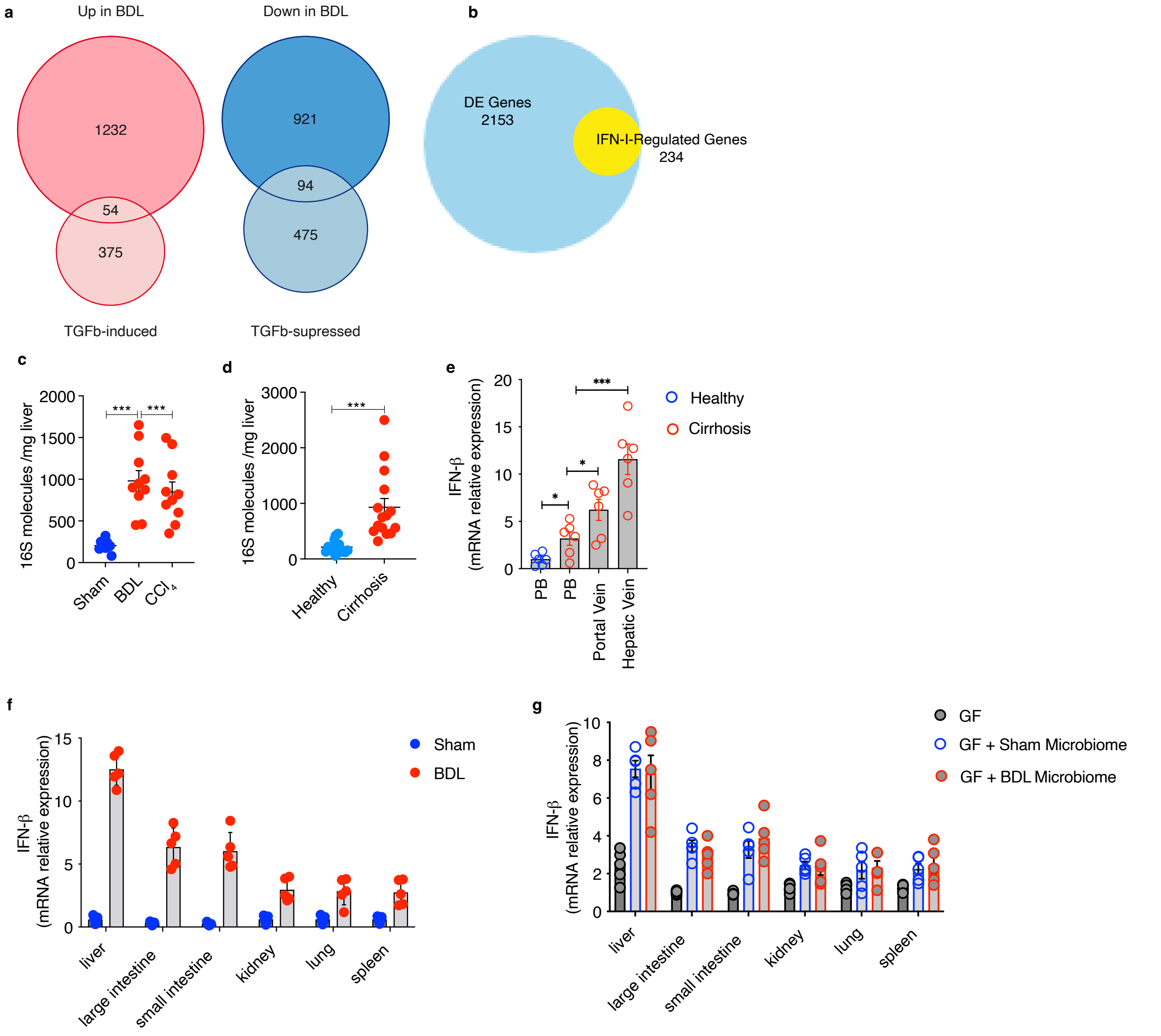


**Extended Data Figure 4 | Bioinformatic analysis of differentially expressed genes in dysfunctional CD8 T cells and intestinal microbiome induced IFN I**

**a**, venn diagrams illustrating overlap between upregulated (left) or downregulated (right) DEGs in LCMV-specific P14 CD8 T cells from liver of BDL-operated mice at d8 p.i. with two gene sets up- or downregulated in response to TGF-β signalling[77]. **b,** Venn diagram illustrating the proportion of IFNAR-related genes among DEGs of P14 CD8 T cells from liver of BDL-operated mice at d8 p.i.. **c,** 16S rDNA quantification in liver of sham or BDL operated (9 days) or $CCl_4$-treated mice (12 weeks). **d,** 16S rDNA quantification in liver tissue biopsies from healthy individuals or cirrhosis patients. **e,** IFNβ mRNA expression in whole blood cells from the portal vein and hepatic vein of cirrhosis patients and from peripheral blood (PB) of healthy subjects or cirrhosis patients. **f**, IFNβ mRNA expression in organs from sham- or BDL-operated mice (day 9). **g**, IFNβ mRNA expression in tissues of BDL-operated Germ Free (GF) mice either untreated or colonized by cecal microbiota from BDL- or sham-operated mice. **b,d-g**, data from two independent experiments; statistics were assessed by unpaired t-tests, **P≤0.01, ***P≤0.001, ****P≤0.001.

Extended Data Figure 5

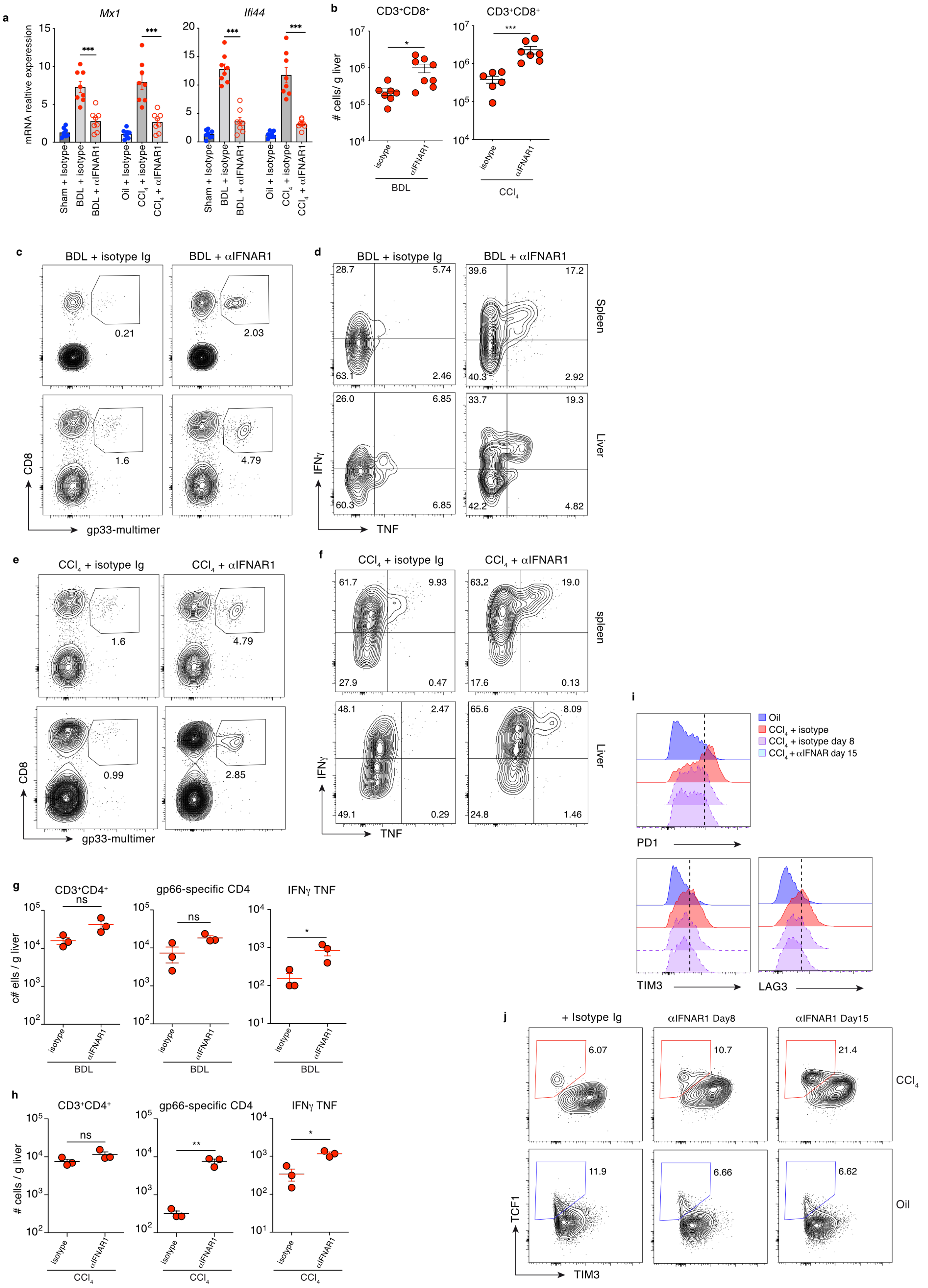

**Extended Data Figure 5 | IFNAR-blockade reconstitutes antiviral T cell immunity during liver injury**

Liver injury was induced in wildtype C57/BL6J mice by BDL operation or $CCl_4$ treatment prior to LCMV infection. Mice were treated with IFNAR1 blocking antibody or isotype control every second day starting at day 5 post infection. **a**, mRNA expression of *Mx1* and *Ifi44* in liver of BDL- or sham-operated and $CCl_4$ or oil-treated mice at d4 p.i. **b,** numbers of CD8 T cells in liver of BDL-mice and $CCl_4$-treated mice after anti-IFNAR antibody treatment. **c,d**, frequencies of LCMV gp33-specific CD8 T cells and their cytokine production in liver of BDL-mice at d8 p.i. **e,f**, frequencies of LCMV gp33-specific CD8 T cells and their cytokine production in liver of $CCl_4$-treated mice at d8 p.i. **g,h,** numbers of CD4, LCMV gp66-specific CD4 and IFNγ / TNF LCMV-specific CD4 T cells in liver of BDL-mice or $CCl_4$-treated mice at d8 p.i. **i,j** expression of co-inhibitory on LCMV gp33-specific $CD8^+$ T cells $CCl_4$-treated mice after anti-IFNAR antibody treatment. Statistics were assessed by unpaired t-tests, ns = not significant, $^*P \leq 0.05$, $^{**}P \leq 0.01$, $^{***}P \leq 0.001$.

Extended Data Figure 6

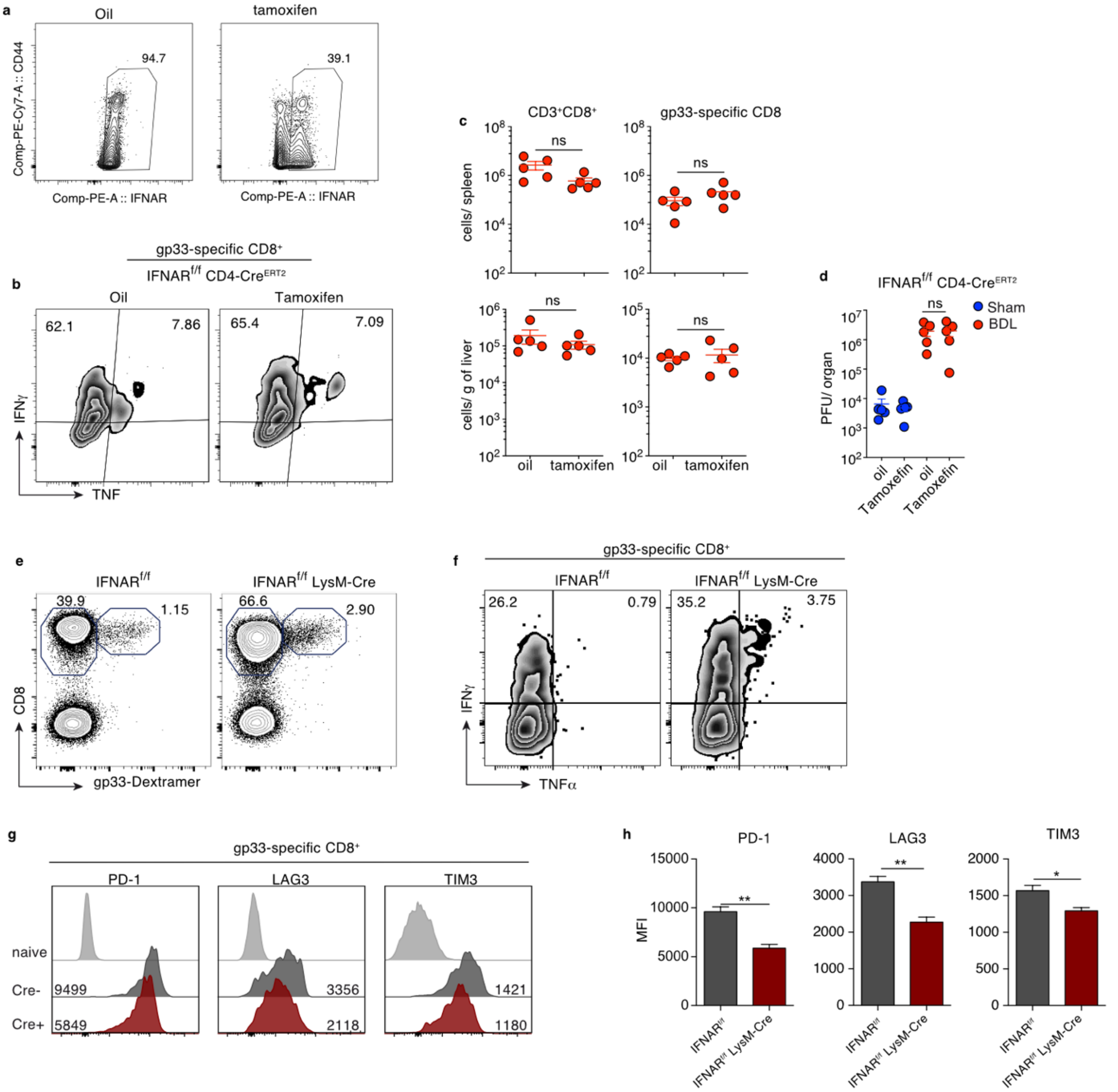


**Extended Data F igure 6 | Myeloid cell-specific IFNAR-blockade reconstitutes anti-viral T cell immunity during liver injury**

**a,** IFNAR1 expression by CD4 T cells from spleen of IFNAR$^{ff}$ x CD4-Cre$^{ERT2}$ mice treated with tamoxifen or oil as control. **b-d**, IFNAR$^{ff}$ x CD4-Cre$^{ERT2}$ BDL-mice received tamoxifen or oil as control and were LCMV infected on day 9 post operation. **b,c,** frequency and numbers of LCMV gp33-specific CD8 T cells and IFNγ– and TNF-producing T cells in spleen at d8 p.i. **d**, LCMV titers in liver at d8 p.i. **e**,**f,** frequencies of LCMV gp33-specific CD8 T cells and their IFNγ and TNF production from the spleen of IFNAR$^{f/f}$ LysM-Cre$^{+}$ mice or Cre$^{negative}$ littermates at d8 p.i. **g**,**h**, expression of co-inhibitory molecules by LCMV gp33-specific CD8 T cells from liver of IFNAR$^{f/f}$ LysM-Cre$^{+}$ mice or Cre-$^{negative}$ littermate BDL-mice at d8 p.i. Statistics were assessed by unpaired t-tests; ns = not significant, *P≤0.05, **P≤0.01

Extended Data Fig. 7

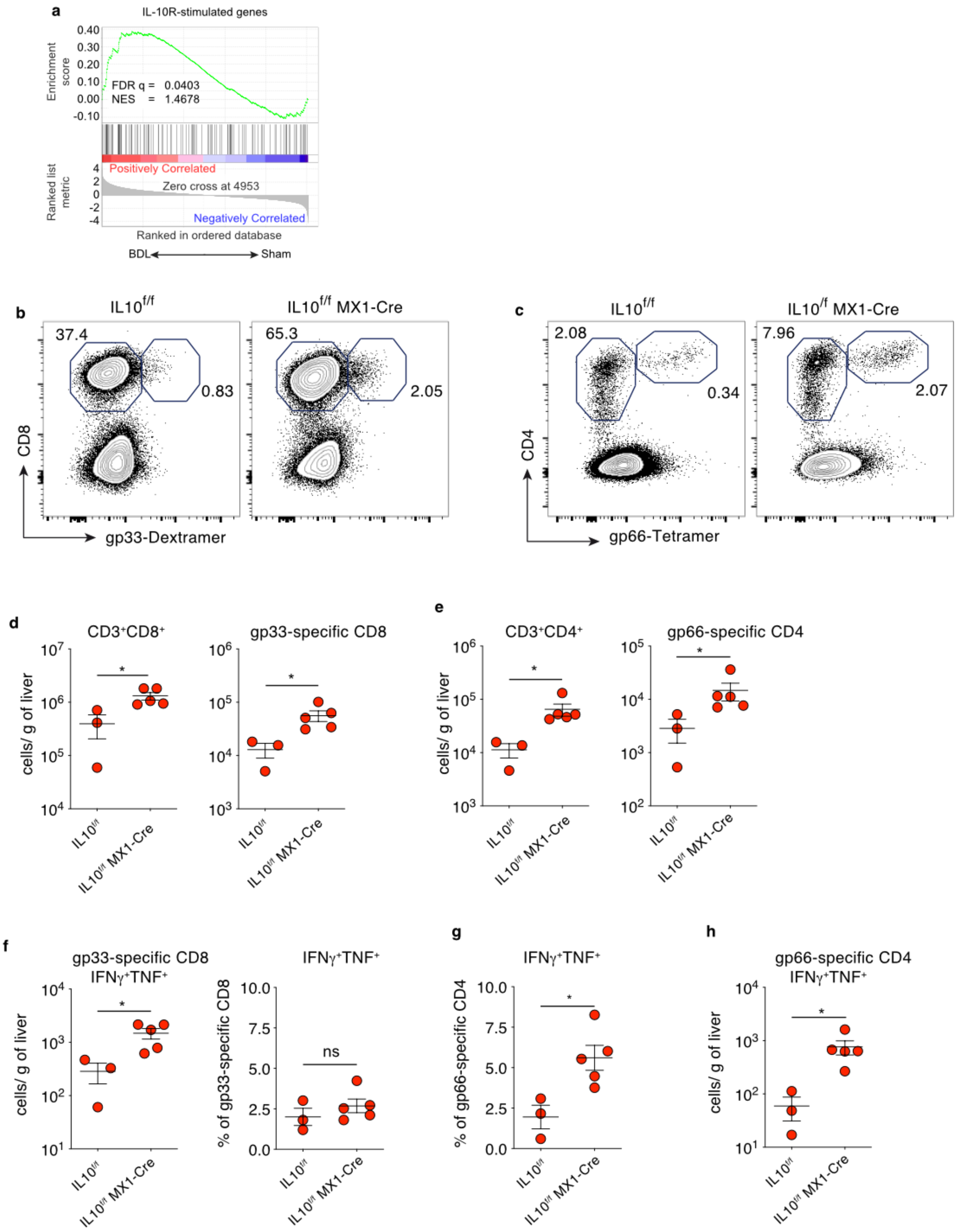

**Extended Data Figure 7 | IFN I-induced IL-10 in suppression of antiviral T cell immunity**

**a,** Gene Set Enrichment analyses (GSEA) depicting enrichment of IL-10Rα-signaling associated gene sets within the transcriptome of LCMV-specific P14 CD8 T cells from BDL-mice at d8 p.i.. **b-h,** liver injury was induced in $IL10^{f/f}$ $Mx1\text{-}Cre^{+}$ mice or $Cre\text{-}^{negative}$ littermates by BDL. **b,d,f,** frequencies and numbers of hepatic $CD8^{+}$ T cells, LCMV gp33-specific CD8 T cells and their IFNγ/TNF production from $IL10^{f/f}$ $Mx1\text{-}Cre^{+}$ mice or $Cre\text{-}^{negative}$ littermate BDL-mice. **c,e,g,h** frequencies and numbers of hepatic CD4 T cells, LCMV gp66-specific CD4 T cells and their IFNγ/TNF production from $IL10^{f/f}$ $Mx1\text{-}Cre^{+}$ mice or $Cre\text{-}^{negative}$ littermate BDL-mice. **b-e**. Data were obtained from two independent experiments Statistics were assessed by unpaired t-tests. ns = not significant, *P≤0.05.

Extended Data Fig 8

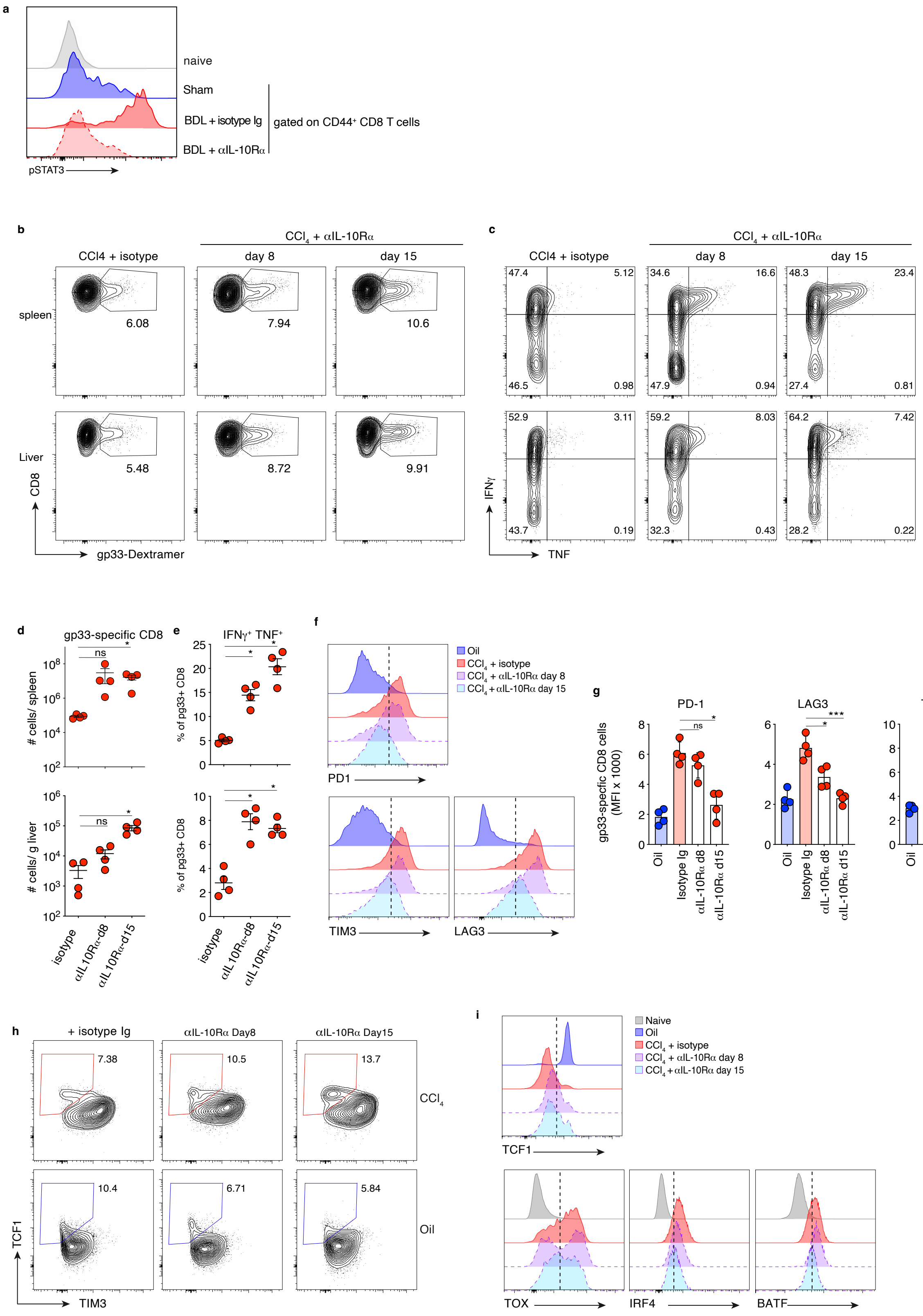

**Extended Data Figure 8 | Blockade of IL-10Rα-signaling restores antiviral CD8 T cell immunity during liver injury**

**a,** phosphorylated STAT3 levels in LCMV gp33-specific CD8 T cells BDL-mice treated with IL10R blocking antibody or isotype control at d2 p.i. **b-e,** frequencies of liver and spleen $CD8^+$ T cells, LCMV gp33-specific CD8 T cells and their IFNγ/TNF-production in $CCL_4$-treated mice after anti-IL-10Rα antibody treatment at d8 and d15 p.i. **f-i**, expression of co-inhibitory molecules and transcription factors by LCMV gp33-specific CD8 T cells in in $CCL_4$-treated mice after anti-IL-10Rα antibody treatment. Data are representative of three independent experiments. Statistics were assessed by unpaired t-tests. ns not significant, *P≤0.05, **P≤0.01. *P≤0.05, **P≤0.01, ***P≤0.001.

Extended Data Fig 9

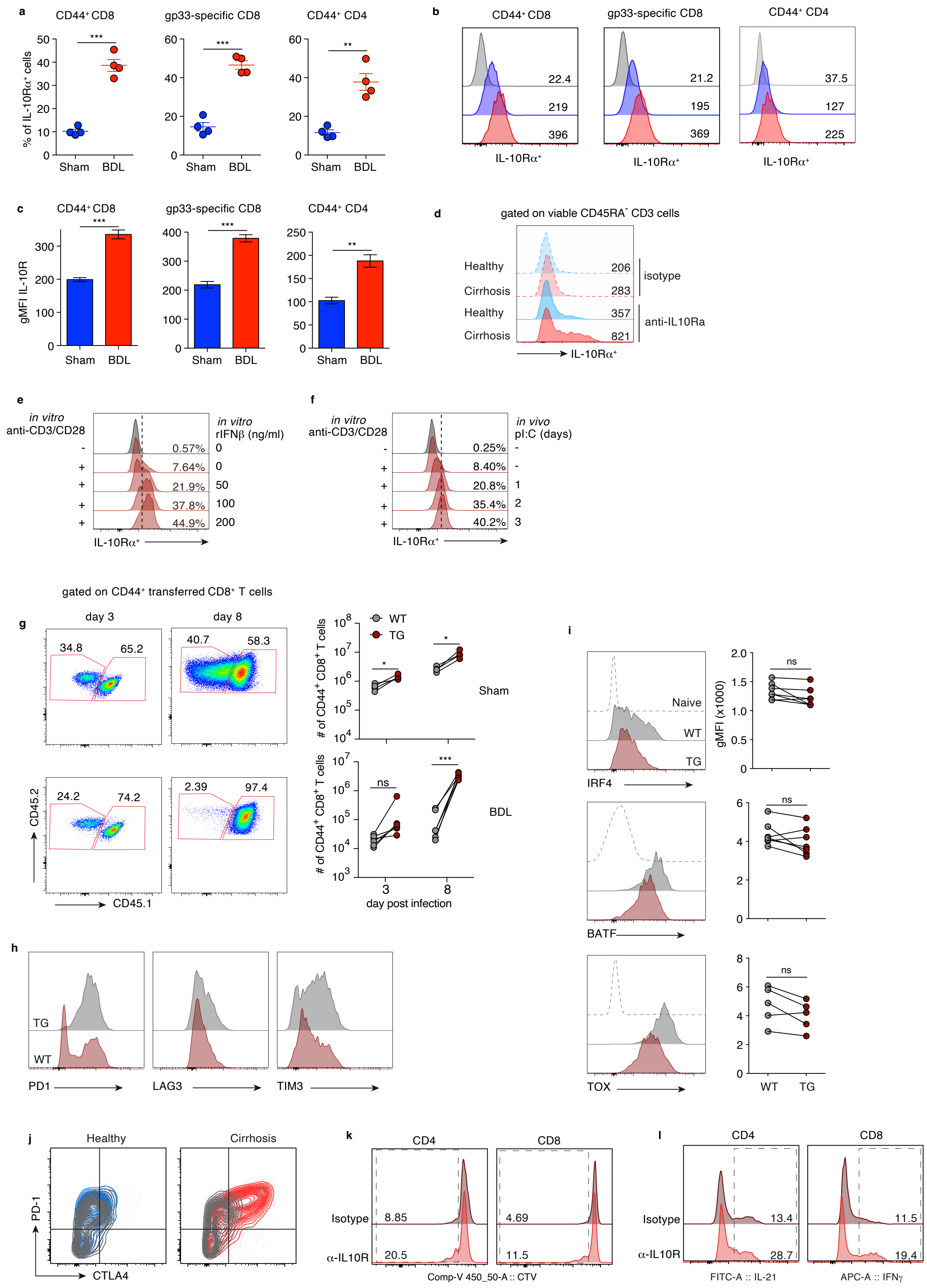

**Extended Data Figure 9 | T cell-specific IL-10Rα-signaling hampers effector functions and blockade of IL-10Rα-signaling in vaccination-induced human T cells restores functionality**

**a-c,** percentage of IL-10Rα expressing immune cells from BDL- or sham-operated mice at d2 p.i. **d**, IL10Rα expression on peripheral blood $CD3^{+}CD45RA^{neg}$ T cells of healthy donors or cirrhotic patients. **e**, IL10Rα expression of murine T cells stimulated *ex vivo* with αCD3/CD28 antibodies in presence of increasing concentrations of IFNβ. **f**, IL10Rα expression after *ex vivo* stimulation with αCD3/CD28 antibodies by mouse T cells isolated at d3 after pI:C application. **g-i**, sub-lethally irradiated wildtype mice ($CD45.1^{+}$) were grafted with 1:1 mixture of naive wildtype (WT, $CD45.2^{+}$) and CD4-dominant negative (DN) IL-10R transgenic (TG, $CD45.1/2^{+}$) T cells. 4 weeks later mice underwent either sham or BDL operation and infected 9 days later with LCMV. **g,** percentage of WT or TG $CD44^{+}$ CD8 T cells in spleen of recipient mice. **h,i**, co-inhibitory molecule expression by LCMV gp33-specific CD8 T cells in liver at d8 p.i. **j**, PD1 and CTLA4 expression by $CD3^{+}$ T cells from healthy or cirrhotic subjects after *ex vivo* stimulation with anti-CD3/CD28 in presence of IL-10Rα blocking (blue and red) or isotype control (grey) antibodies. **k,i**, dilution of CTV dye (**k**) or expression of IL-21 or IFNγ (**l**) by HBs-specific CD4 and CD8 T cells from healthy (n=4) or cirrhotic (n=4) donors after ex vivo peptide-specific stimulation in presence of IL-10Rα blocking antibodies. Data representative of ≥2 independent experiments. **a,c,** Statistics were assessed by unpaired t-tests. *P≤0.05, **P≤0.01, ***P≤0.001. Comparison between groups was calculated using a two-tailed paired Student's t-test, ns not significant, **P≤0.01, ***≤P0.001.
